# Adoptive cell therapy using T cell receptors equipped with ICOS yields durable anti-tumor response

**DOI:** 10.1101/2025.10.14.682056

**Authors:** Alexandre Marraffa, Cor Berrevoets, Margherita Mosiello, Rebecca Wijers, Daphne Roelofs, Mandy van Brakel, Kim Kroese, Marlies J.W. Peeters, Willem A. Dik, Andre Kunert, Rachel J.M. Abbott, Dora Hammerl, Christopher Schliehe, Reno Debets

**Affiliations:** Laboratory of Tumor Immunology, Department of Medical Oncology, Erasmus MC, Rotterdam, the Netherlands; Pan Cancer T BV, Rotterdam, the Netherlands; Laboratory of Medical Immunology, Department of Immunology, Erasmus MC, Rotterdam, the Netherlands; Department of Immunology, Erasmus MC, Rotterdam, the Netherlands

**Keywords:** co-stimulatory TCR-T cell, immunotherapy, tumor immunology, gene engineering, solid tumors

## Abstract

Treatment with adoptively transferred T cells is challenged by limited longevity of therapeutic cells within tumors. To enhance the durability of anti-tumor T cell products, we have created T cell receptors (TCRs) with built-in co-stimulatory molecules. We observed that TCRs coupled to ICOS mediated exceptionally long-term responses including delay of tumor recurrence and cures in a mouse tumor model. TCR:ICOS T cells showed enhanced and antigen-specific production of inflammatory cytokines and resistance to exhaustion. Genetic ablation of ICOS-mediated activation of the PI3K-NFκB pathway neutralized the long-term anti-tumor effects. To translate TCR:ICOS to human T cells, we identified a single amino acid change in the cytosolic tail which was necessary for functional surface expression. Notably, the optimized receptor sustained performance of human T cells upon repeated stimulation across multiple antigen specificities. Collectively, we present a novel and uniformly applicable TCR:ICOS format that supports fitter T cell products for adoptive cell therapy.

**Highlights:** Newly designed co-stimulatory TCR, with extracellular TCR-V and C domains coupled to CD28 transmembrane domain, and ICOS and CD3ε intracellular domains (in short TCR:ICOS) provides:

➢ durable anti-tumor response and T cell persistence in mouse model
➢ inflammatory T cell phenotype and resistance to T cell exhaustion
➢ effects via PI3K and NFκB activation
➢ translation to human T cells upon single amino acid mutation in TCR:ICOS tail
➢ extension to multiple clinically relevant TCRs while preserving prolonged T cell fitness

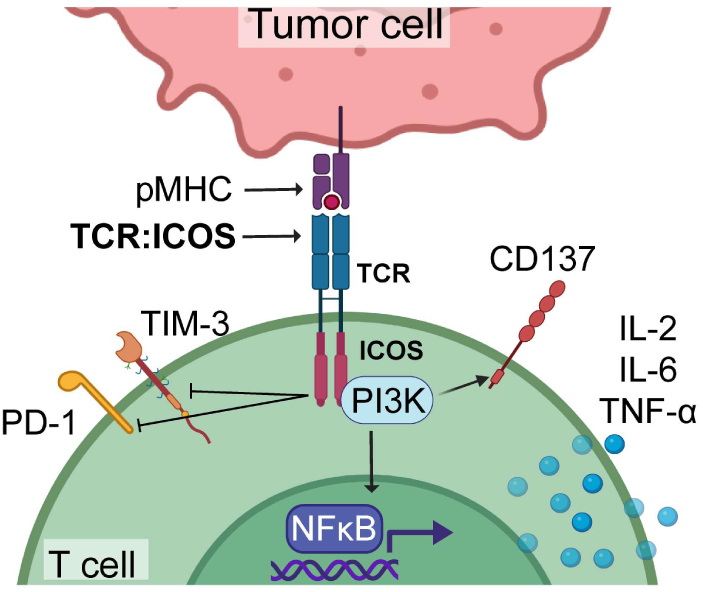

## Introduction

Adoptive cell therapy to treat cancer relies on the infusion of tumor-specific T cells that subsequently trace and kill tumor cells. Besides non-modified T cells, such as tumor-infiltrating lymphocytes (TILs) that are directed against cancer antigens via endogenous T cell receptors (TCRs), T cells are often derived from blood and redirected toward cancer antigens via gene-engineering with chimeric antigen receptors (CARs) or TCRs. CAR-T cell therapy has already exhibited remarkable success and achieved objective response rates up to 90% in hematological malignancies (1–6). Currently, seven CAR-T cell products directed to either CD19 or BCMA have received FDA approval and now constitute standard-of-care treatment for B cell lymphomas and multiple myeloma (7). In addition, TIL and TCR-T cell treatments have shown objective responses of over 40% in solid tumors, such as melanoma and synovial sarcoma (8,9). This has led to the 2024 FDA approval of a TIL product directed against melanoma and a TCR-T cell product directed against synovial sarcoma antigen MAGE-A4 (10). Despite this tremendous progress, there is still an urgent need to identify gene-engineering strategies to enhance T cell fitness, particularly for the treatment of solid tumors, where the long-term efficacy of T cell therapy is lagging behind that of hematological malignancies.

A major cause of ineffective or transient clinical effects is the inability of adoptively transferred T cells to sufficiently resist the immunosuppressive tumor microenvironment (TME). Multiple reports have already demonstrated that within TMEs, T cells become progressively dysfunctional, show a curtailed persistence as well as a diminished anti-tumor activity, as reviewed in (11–13). In addition, the inhospitable TME impairs T cell infiltration, migration and antigen presentation to T cells, which, together, significantly limit the durability of anti-tumor T cells (12). Specifically, TMEs lack sufficient co-stimulatory ligands, thereby preventing optimal T cell activation and potentially initiating T cell impairments (14,15). In fact, negligible expression of co-stimulatory ligands, often accompanied by an overabundance of co-inhibitory ligands, may mirror TMEs that are the result of adaptive feedback mechanisms that hamper the longevity of anti-tumor T cells (16–19). To bypass the need for exogenous, i.e. TME-derived co-stimulation, we propose to create TCRs with built-in co-stimulatory molecules, thereby providing a second signal upon antigen recognition. Although clinical CARs already contain co-stimulatory domains, engineering co-stimulatory TCRs with the potential to unlock T cell therapy for a large collection of targets, is an underexplored strategy. In fact, engineering TCRs, in contrast to CARs, faces typical structural constraints due to the necessary assembly of TCR chains with CD3 components as well as the requirement to bind peptide:MHC complexes.

In the present study, we have developed a panel of new TCRs incorporating co-stimulatory molecules and evaluated these TCR-T cells for durability of the anti-tumor effect in an aggressive melanoma mouse model. The TCR harboring Inducible T cell Co-Stimulator (TCR:ICOS) resulted in exceptionally long-term cures and delay of tumor recurrence and was identified as the lead candidate. Subsequently, we uncovered its mechanism of action by performing a series of *in vivo* and *in vitro* experiments. Finally, we translated the murine TCR:ICOS format for its use in human T cells and demonstrated its surplus value regarding T cell longevity across multiple clinically relevant antigen specificities.

## Materials and Methods

### Design of murine co-stimulatory TCRs

TCRα and β genes specific for the human gp100_280-288_ epitope (YLEPGPVTA) presented by HLA-A2 (gp100/A2) were derived from clone CTL-296 (20), murinized and codon-optimized as described earlier (21). Subsequently, TCR genes were cloned into the pMP71 vector (a kind gift of Prof. Wolfgang Uckert, MDC, Berlin, Germany (22)), in which the TCRα and β chains were separated by an optimized T2A ribosome skipping sequence according to a TCRß-T2A-TCRα format, as described before (23). Murine TCR:CD28 was generated as described in Govers et al., 2014 (24). Shortly, human (h)TCR-Vα was coupled to mouse (m)TCR-Cα and hTCR-Vß to mTCR-Cß (all extracellular domains), and each TCR chain was followed by a proline-lysine (PK) linker, the mCD28 transmembrane, mCD28 intracellular and mCD3ε intracellular domains. Murine OX40, ICOS and 4-1BB intracellular domains were ordered via GeneArt (Life Technologies) and introduced into the TCR:CD28 cassette (replacing the intracellular domain of CD28) via overlap PCR with use of Q5 High Fidelity DNA Polymerase (New England Biolabs). One variant to TCR:ICOS with an amino acid substitution in ICOS (Y181F), named TCR:ICOS-YF, was designed by site directed mutagenesis of Y181F in the mICOS domain to prevent docking and downstream signaling of PI3K, as described before by Gigoux et al. (25). Sequences were obtained and checked using the BigDye Terminator v3.1 Cycle Sequencing Kit (Applied Biosystems) and the ABI sequence analyzer. All mTCRs used in this study are schematically depicted in **Figures 1a** and **4a**. Primer sequences can be obtained upon request.

**Figure 1.**
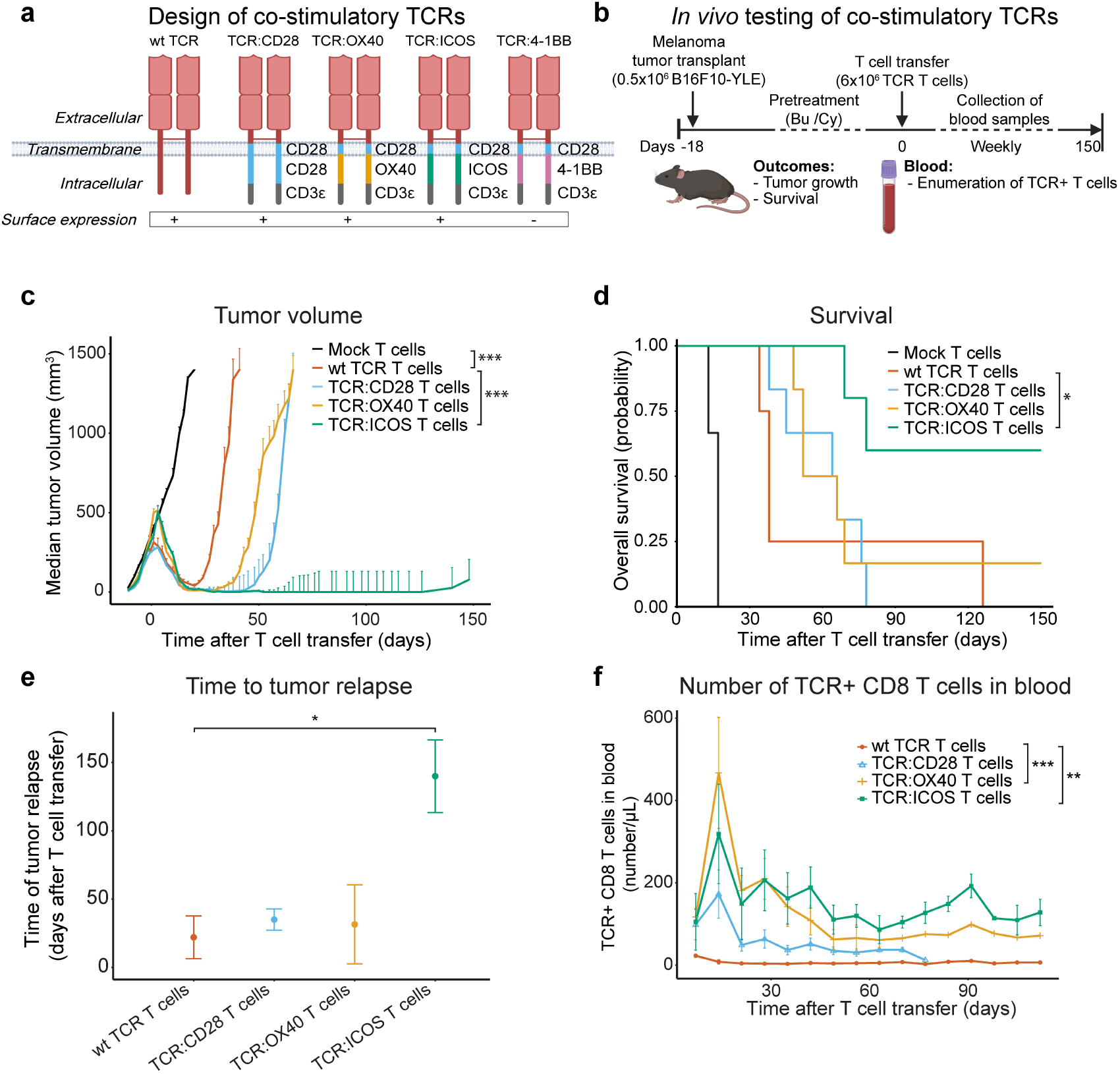
TCR:ICOS mediates durable anti-melanoma T cell responses. **(a)** Schematic illustration of the design of murine co-stimulatory TCRs, where extracellular TCR-VαCα and VßCß chains targeting human gp100/A2 were coupled to the murine transmembrane domain of CD28, followed by the murine intracellular domains of either CD28, OX40, ICOS or 4-1BB, and the intracellular domain of CD3ε. **(b)** C57BL/6 HHD mice were transplanted with B16:HHD-YLE cells expressing the HLA-A2-restricted gp100 epitope YLEPGPVTA. Prior to adoptive T cell transfer on day 0, mice received busulfan and cyclophosphamide, after which mice were administered 6×10^6^ mock, wt TCR, TCR:CD28, TCR:OX40 or TCR:ICOS T cells. Mice were then monitored for 150 days after T cell transfer for tumor volume, survival and number of TCR T cells in blood. **(c)** Tumor volume, displayed in mm^3^ following T cell transfer (median+SEM, n=4-6 per group). Mice attaining a tumor volume>1400 mm^3^ were sacrificed. Statistical testing between groups was performed with linear mixed model with elapsed time as a separate variable and correction for random effect. Significant interactions between time and treatment on tumor volume is displayed. **(d)** Survival of mice showing initial response following T cell transfer is plotted as Kaplan-Meier curves. Statistical inter-group comparison was done according to Cox regression. **(e)** Time to tumor relapse in days following T cell transfer is plotted (median±SEM, n=4-6). Inter-group comparison was done according to Kruskal-Wallis testing, with post-hoc Dunn’s test. **(f)** Numbers of TCR+ CD8 T cells per µL blood following T cell transfer are displayed as determined by flow cytometry (median±SEM, n=4-6). Inter-group comparison was done according to linear mixed model with elapsed time as a separate variable and correction for random effect. With * = p<0.05, ** = p<0.01, *** = p<0.001.

**Figure 2.**
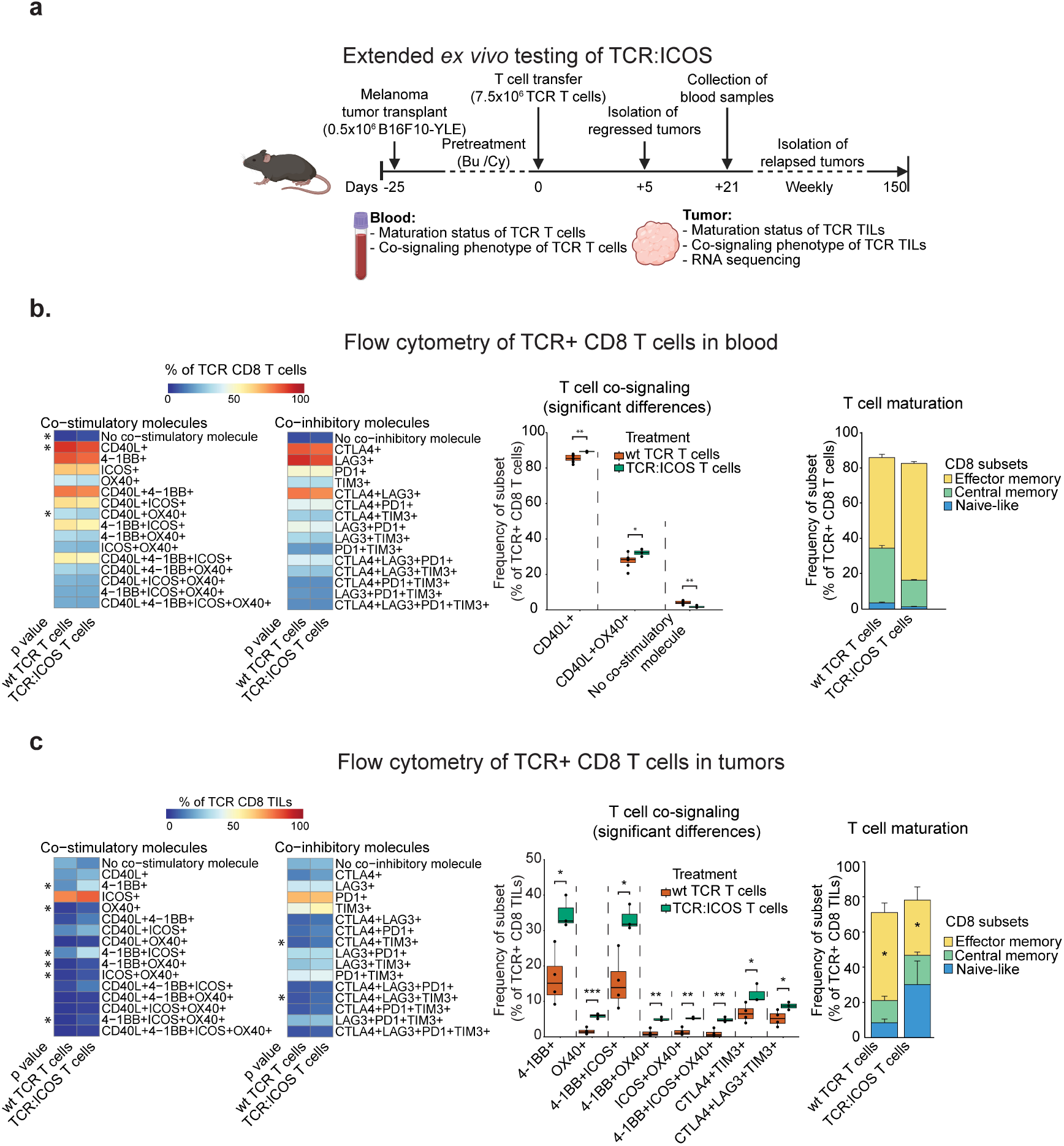
Intra-tumoral TCR:ICOS CD8 T cells exhibit a young and co-stimulatory phenotype. **(a)** Experimental design, in which C57BL/6 HHD mice were transplanted with B16:HHD-YLE melanoma cells on day-25 and pre-treated with busulfan and cyclophosphamide, after which mice were administered 7.5×10^6^ wt TCR or TCR:ICOS T cells on day 0. Monitoring included frequency assessment of subsets of TCR+ CD8 T cells in blood (at three weeks after T cell transfer) and TILs (in regressing tumors at five days after T cell transfer) according to maturation and co-signaling status as well as transcriptomics of tumors **(b)** Left panel: heatmap corresponding to phenotype of TCR+ CD8 T cells in blood as determined by flow cytometry. Co-signaling phenotype of T cells was determined according to the expression of (combinations of) co-inhibitory (CTLA4, LAG3, PD1, TIM3) or co-stimulatory receptors (CD40L, 4-1BB, ICOS, OX40). The heatmap displays median expressions of single or multiple markers, n=3-8 per group. Statistical significance between TCR:ICOS and wt TCR T cells was assessed according to t-test and is highlighted at left-hand side. Middle panel: TCR:ICOS+ CD8 T cell subsets that show differential frequencies when compared to wt TCR+ CD8 T cells. Right panel: stacked bars corresponding to maturation phenotype of TCR+ CD8 T cells in blood as determined by flow cytometry (n=3-8 per group). Maturation status of T cells was defined as follows: naïve-like T cells, CD62L+CD44-; central memory T cells, CD62L+CD44+; and effector memory T cells, CD62L-CD44+. Statistical significance between TCR:ICOS and wt TCR T cells was assessed according to Kruskal-Wallis testing. **(c)** Left and right panels: heatmap and stacked bars corresponding to phenotype of TCR+ CD8 TILs from regressing tumors as determined by flow cytometry. Middle panel: TCR:ICOS+ CD8 TIL subsets that show differential frequencies when compared to wt TCR+ CD8 TILs (n=2-4 per group). With *= p<0.05, ** = p<0.01, *** = p<0.001.

**Figure 3.**
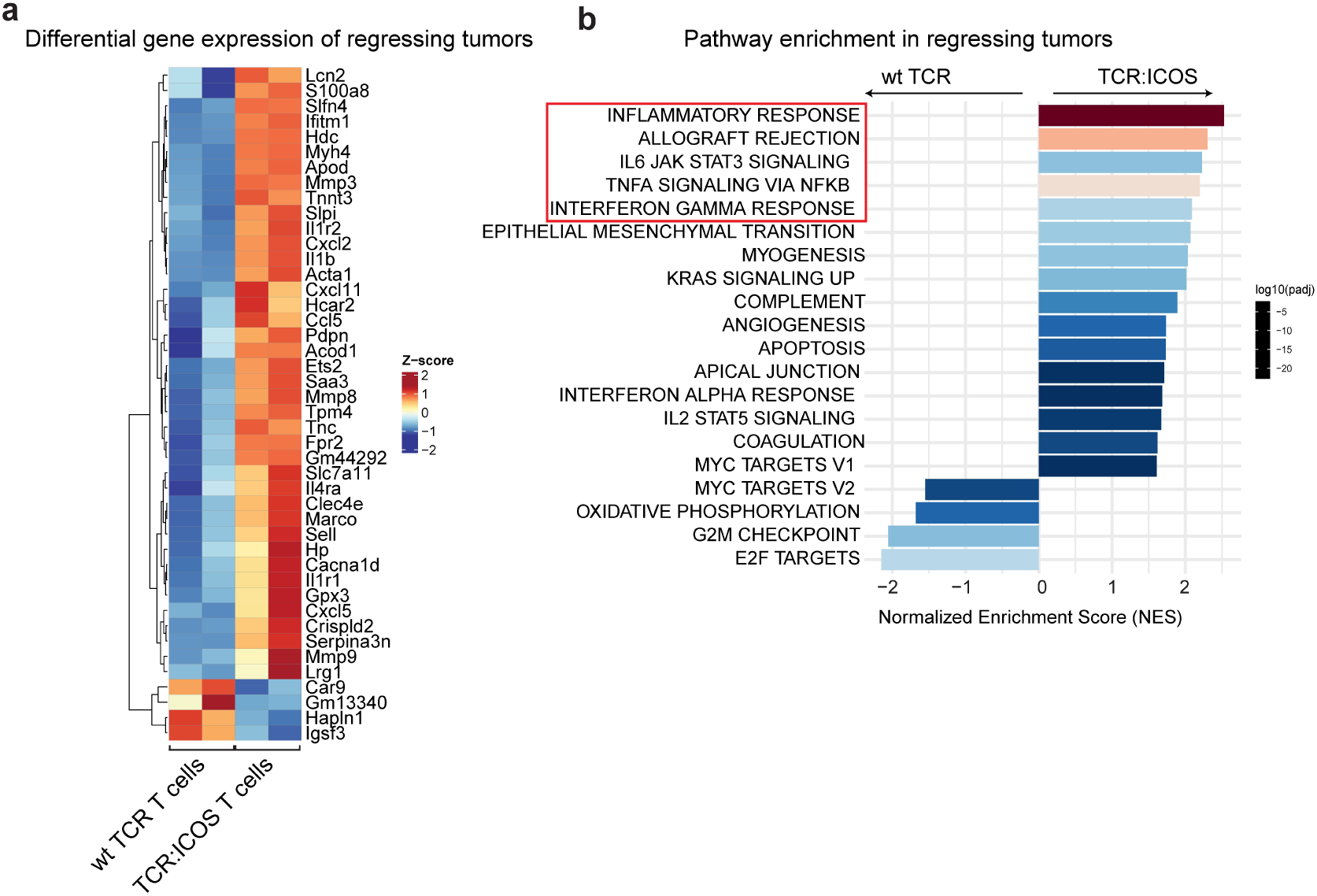
TCR:ICOS T cells create an inflammatory micro-environment in tumors. Regressing tumor tissues from mice treated with wt TCR T cells or TCR:ICOS T cells were isolated on day five for transcriptomic analysis. **(a)** Heatmap displaying all genes with an absolute log2FC >2, a base mean >50 and an adjusted p-value<0.05 for TCR:ICOS T cells when compared to wt TCR T cells (n=2 per group). **(b)** Bar graph showing top 20 gene sets that are differentially enriched following transfer of TCR:ICOS T cells or wt TCR T cells.

**Figure 4.**
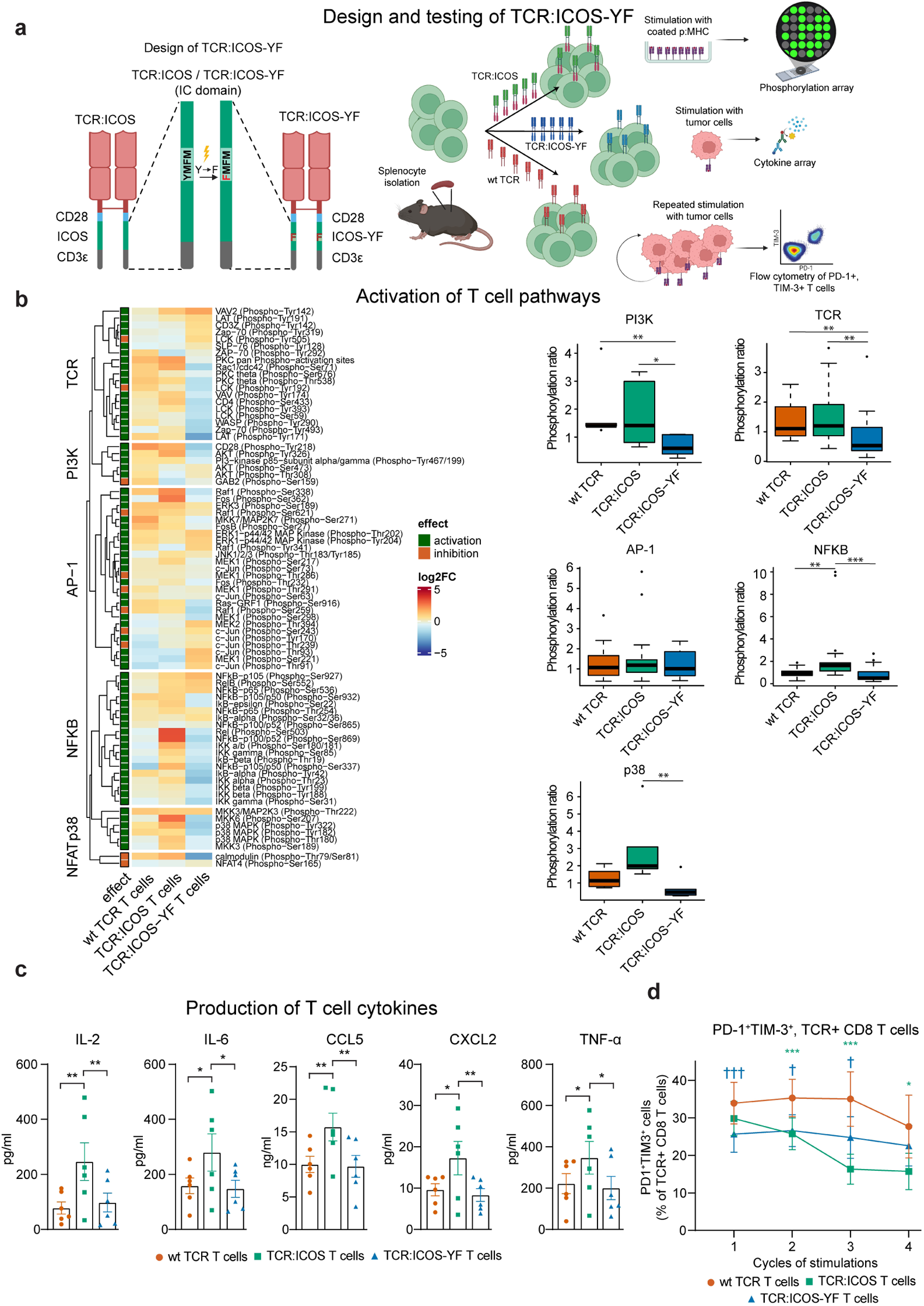
In vitro stimulated TCR:ICOS T cells activate the NFκB pathway, produce inflammatory cytokines and are resistant to exhaustion upon repeated tumor cell exposure. **(a)** Left panel: graphical representation of TCR:ICOS-YF, where the intracellular ICOS domain is mutated at a single site to prevent docking of PI3K and downstream signaling. Right panel: experimental design in which mouse splenocytes that express either TCR:ICOS, TCR:ICOS-YF or wt TCR were interrogated for intracellular signaling, cytokine production and ability to preserve fitness upon repeated antigen stimulation. **(b)** Left panel: heatmap corresponding to fold-change in protein phosphorylation in wt TCR, TCR:ICOS and TCR:ICOS-YF T cells upon stimulation. Mouse T cells were incubated for 30 min at 37°C with pre-coated A2Kb pentamers presenting the cognate epitope, after which cells were lysed and lysates were exposed to a fluorescence-based phosphorylation array covering more than 70 substrates representing T cell signaling. In the heatmap, phosphorylated targets were categorized according to different intracellular pathways. The effect of phosphorylation on a particular pathway is displayed to the left, with red representing an inhibitory and green an activating substrate. Right panel: box plots displaying ratio of phosphorylated substrates associated with different pathways, only taking into account pathway-activating substrates. Comparison between groups was performed using a Kruskal-Wallis, followed by post-hoc Dunn’s test. **(c)** Bar graphs displaying cytokine production of TCR T cells following stimulation with tumor cells. T cells were incubated for 16h at 37°C with B16 cells that expressed the cognate epitope, after which supernatants were assayed for the presence of 14 cytokines according to a cytokine bead array. Statistical significance between groups was tested with two-way ANOVAs with Dunnet’s multiple comparison test. Only the conditions in which significant differences were observed are displayed here, with the full panel of cytokines being available in **Supplementary Figure 4c. (d)** Average percentages of PD1^+^TIM3^+^ TCR+ CD8 T cells following repeated stimulations (±SEM, n=9). Co-culture with B16F10 wt tumor cells not expressing the cognate epitope did not result in changed percentages of this T cell subset and was used to normalize the background expression. Statistical significance between groups was tested with a mixed model analysis, and post-hoc analysis per cycle was performed with Dunnet’s multiple comparison test. Statistical significance between TCR:ICOS and wt TCR T cells is depicted with a star (*), and between TCR:ICOS-YF and wt TCR with a cross (✝). With * = p<0.05, ** = p<0.01, *** = p<0.001.

### Design of human co-stimulatory TCRs

To convert the gp100/A2 mTCR:ICOS into a human format (hTCR:ICOS), we generated the pMP71 vector consisting of gp100/A2 hTCR-VDJß-Cß-T2A-hTCR-VJα-Cα (extracellular domains), and in which each TCR chain was followed by a proline-lysine (PK) linker, the hCD28 transmembrane, the hICOS intracellular and hCD3ε intracellular domains. This hTCR:ICOS construct demonstrated poor surface expression in human T cells (see **Supplementary Figure 6a**). To identify which building blocks (or combinations thereof) and/or which amino acid boundaries of such building blocks determine functional expression of TCR:ICOS in human T cells, a panel of 49 variants was designed (all produced by GeneArt), and tested for surface expression in a stepwise approach. An overview of the different constructs, including detailed specifics, is provided in **Supplementary Table 1**. In all transduction experiments, the wt TCR was taken along as a reference. A scheme of the study design to select and test hTCR:ICOS variants is shown in **Figure 6a**. Following selection of hTCR:ICOS-RA, in which the mutation R54A was introduced in the hCD3ε domain (**Figure 6c)**, this construct was further tested for antigen recognition, sensitivity, cytotoxicity and performance upon repeated antigen stimulation. To test the applicability of hTCR:ICOS-RA to other TCR specificities, the gp100/A2 TCR-Vα and Vß domains were exchanged with those of the TCR directed against NY-ESO1/A2 and against Ropporin-1/A2 (ROPN1, all new constructs were produced by GeneArt) (26–28). The NY-ESO1 hTCR:ICOS-RA and the ROPN1 hTCR:ICOS-RA were tested again for surface expression, antigen recognition, sensitivity and performance upon repeated antigen stimulation.

**Figure 5.**
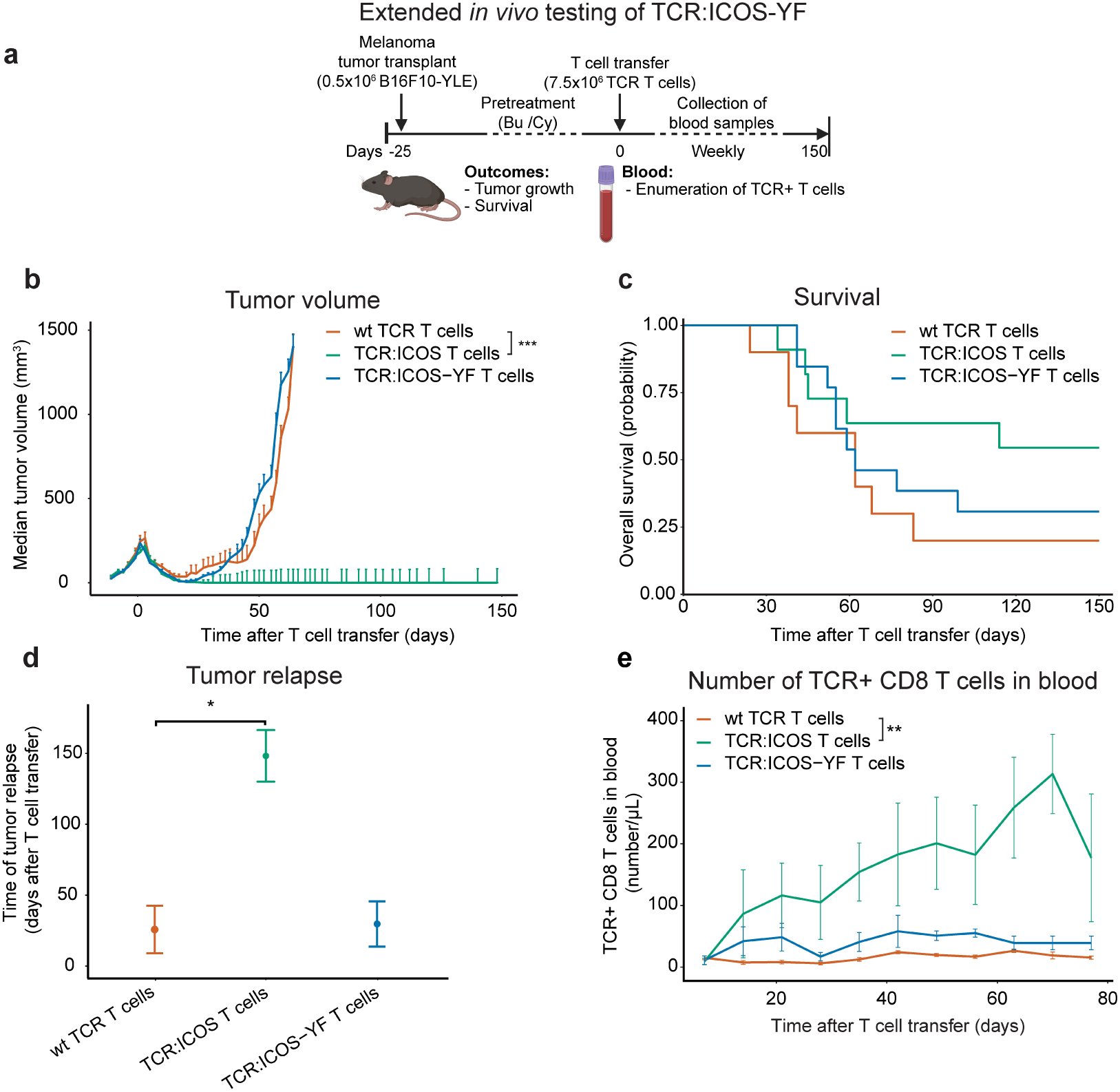
Durability of anti-tumor response of T cells depends on ICOS docking site for PI3K. **(a)** TCR:ICOS-YF T cells were studied for their *in vivo* anti-tumor performance. Schematic illustration of experimental design, in which mice were transplanted with B16:HHD-YLE melanoma cells and pre-treated with chemotherapy, after which mice were administered 7.5×10^6^ wt TCR, TCR:ICOS or TCR:ICOS-YF T cells. Monitoring included tumor growth and assessment of subsets of TCR+ CD8 T cells in blood. **(b)** Line graph displaying tumor volume in mm^3^ following T cell transfer (median+SEM, n=10-13 per group, pooled from 2 independent experiments). Statistical testing between groups was performed with linear mixed model with elapsed time as a separate variable and correction for random effect. Significant interaction between time and treatment on tumor volume is displayed. **(c)** Survival of mice following initial response to T cell transfer is plotted as Kaplan-Meier curves. **(d)** Time to tumor relapse in days following T cell transfer is plotted (median±SEM, n=10-13, pooled from 2 independent experiments). Inter-group comparison was performed using a Kruskal-Wallis test. **(e)** Number of TCR+ CD8 T cells per µL blood following T cell transfer is plotted as determined by flow cytometry (median±SEM, n=10-13, pooled from 2 independent experiments. Statistical testing between groups was performed with linear mixed model with elapsed time as a separate variable and correction for random effect. Significant interaction between time and treatment on T cell number is displayed. With * = p<0.05, ** = p<0.01, *** = p<0.001.

**Figure 6.**
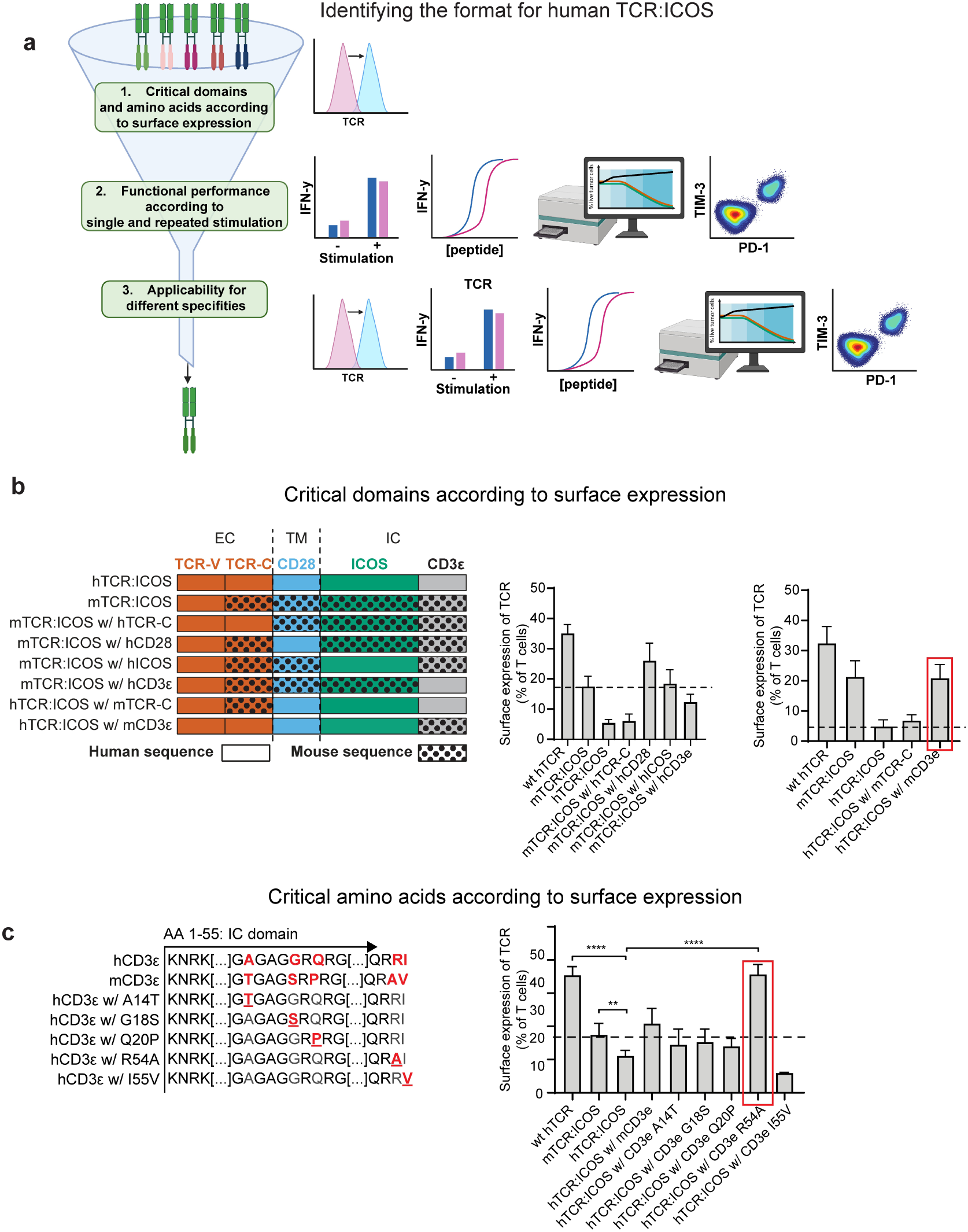
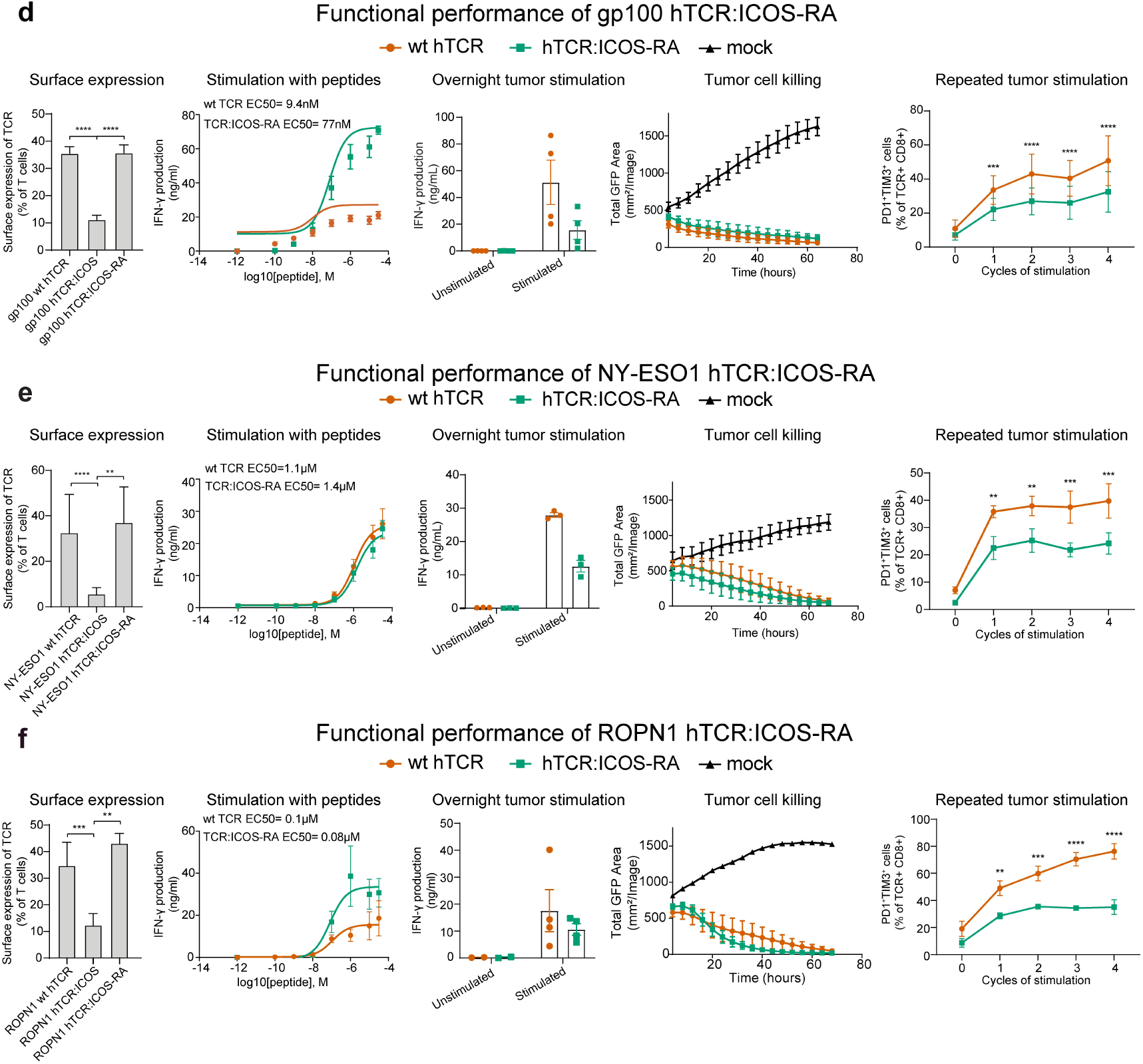
A single amino acid substitution in the cytosolic tail of the CD3ε domain enables functional expression of TCR:ICOS in human T cells. **(a)** Schematic illustration of the selection and testing strategy of hTCR:ICOS variants. **(b)** Identification of single building blocks that are required for surface expression of mTCR:ICOS. Left panel: cartoon of the first series of mouse (m) / human (h) TCR:ICOS variants, with mouse sequences highlighted by black dots. Right panels: surface expression of m/h TCR:ICOS variants according to % TCR-Vß14 of TCR T cells corrected for expressions of Mock T cells (mean±SEM, n=6). TCR:ICOS variant with red square was used as basis for further experimentation. **(c)** Identification of single amino acids within mCD3ε that are required for surface expression of hTCR:ICOS. Left plot: amino acid (aa) alignment between human and mouse CD3ε and overview of all single substituent mutants made. Right plot: surface expression of hTCR:ICOS with human to mouse CD3ε aa substitutes according to % TCR-Vß14, after correction for Mock (mean±SEM, n=2-10). Statistical significance between modified TCR:ICOS variants and hTCR:ICOS was assessed using a Student’s t-test. Again, the TCR:ICOS variant with red square was used as basis for further experimentation. **(d)** Demonstration of antigen reactivity, sensitivity and resistance against exhaustion upon repeated stimulation of hTCR:ICOS-RA. 1^st^ plot from the left: bar graph displaying surface expression of gp100 wt hTCR, hTCR:ICOS, and hTCR:ICOS-RA according to % TCR-Vß14 after correction for expression on mock T cells (mean±SEM, n=10). Statistical significance was calculated using a Sidak-Holm’s test. 2^nd^ plot: IFN-γ production following stimulation with T2 cells that were pre-loaded with titrated amounts of human gp100 peptide. TCR T cells were co-cultured with peptide-loaded T2 cells for 16h at 37°C and supernatants were assayed using an IFN-γ ELISA (mean±SEM, n=4). EC50 values, mentioned in the plot, were calculated using graphpad prism. 3^rd^ plot: bar graph displaying IFN-γ production of TCR T cells following stimulation with BLM tumor cells. TCR T cells were co-cultured with BLM cells that either expressed the human gp100 antigen or not for 16h at 37°C, after which supernatants were assayed by ELISA for IFN-γ (mean±SEM, n=4). Statistical significance between hTCR:ICOS and wt TCR was accessed by two-way ANOVA. 4^th^ plot: T cells were co-cultured with BLM tumor cells expressing the human gp100 antigen coupled to eGFP, and viable tumor cells remaining were quantified using Incucyte (mean±SEM, n=3). 5^th^ plot: T cells were repeatedly stimulated overnight with BLM tumor cells expressing the human gp100 antigen, with the percentage of TCR+ CD8 T cells co-expressing PD1 and TIM3 after each cycle being measured by flow cytometry (mean±SEM n=3). Statistical significance between TCR:ICOS and wt TCR T cells was tested using a two-way ANOVA with Sidak’s post-hoc testing. **(e,f)** The hTCR:ICOS format with R54A is extendable to other TCR specificities, as shown for TCRs specific for NY-ESO1/A2 **(e)** and ROPN1/A2 **(f)**. **(e)** 1^st^ plot from the left: bar graph displaying surface expression of wt TCR and TCR:ICOS according to % TCR-Vß13.1 after correction for Mock (mean±SEM, n=6-12). Statistical significance between hTCR:ICOS-RA and wt TCR was assessed according to Sidak-Holm’s. 2^nd^ plot: IFN-γ production following stimulation with T2 cells that were pre-loaded with titrated amounts of peptide (mean±SEM, n=7). 3^rd^ plot: bar graph displaying IFN-γ production following stimulation with human MM-231-NY-ESO1 cells (mean±SEM, n=3). 4^th^ panel: line graph displaying killing of MM-231-NY-ESO1 cells by T cells (mean±SEM, n=3). 5^th^ panel: change in percentage of PD1+, TIM3+ TCR+ CD8 T cells following repeated 24h stimulations with MM-231-NY-ESO1 cells (mean±SEM, n=5). **(f)** 1^st^ plot: bar graph displaying surface expression of wt TCR and TCR:ICOS according to % TCR-Vß13.1 after correction for Mock (mean±SEM, n=4-8). Statistical significance between hTCR:ICOS-RA and wt TCR was assessed according to Sidak-Holm’s. 2^nd^ plot: IFN-γ production following stimulation with T2 cells that were pre-loaded with titrated amounts of peptide (mean±SEM, n=4). 3^rd^ plot: bar graph displaying IFN-γ production following stimulation with human MM-231-ROPN1 cells (mean±SEM, n=2-4). 4^th^ panel: line graph displaying killing of MM-231-ROPN1 cells by T cells (mean±SEM, n=6). 5^th^ panel: change in percentage of PD1+, TIM3+ TCR+ CD8 T cells following repeated 24h stimulations with MM-231-ROPN1 cells (mean±SEM, n=3). With * = p<0.05, ** = p<0.01, *** = p<0.001.

### Retroviral transduction of TCRs into T cells

Mouse T cell transductions were performed as previously described by Pouw and colleagues (21). Shortly, 293T and Ph-A packaging cells were transfected with pHit60 MLV GAG/POL and pHit123 ENV helper constructs (29), and the TCR of interest, using calcium phosphate (Promega Profection Mammalian Transfection system). Mouse splenocytes were isolated, activated for 24h with 0.5µg/mL Concavalin A (Sigma) and exposed once to virus supernatant produced by the packaging cells. Human T cell transductions were performed as previously described by Lamers and colleagues (30). In this case, 293T and Ph-A packaging cells were transfected with pHit60 MLV GAG/POL and pColtGalV ENV helper constructs (31), and the TCR of interest, using the procedure described above. Human PBMCs were activated for 48h using 10 ng/mL soluble anti-CD3 antibody (OKT3, Thermo Fisher Scientific) before being exposed twice to virus supernatant. TCR expression levels were determined using flow cytometric detection of the introduced TCR-Vß, corrected for percentage positivity as observed with mock-transduced T cells from the same donor, and displayed as % positive cells within the CD3+ T cell population. In experiments measuring killing ability of human TCR T cells, a protocol adapted from a Good Manufacturing Practice (GMP) protocol was followed, as described before (32). Shortly, human PBMCs, isolated from buffy coats obtained from Sanquin (Amsterdam, the Netherlands), were activated with 30 ng/mL soluble anti-CD3 (OKT3, Miltenyi Biotec) and 30 ng/mL anti-CD28 (clone 15E8, Miltenyi Biotec), 110 IU/mL hIL15 (Miltenyi Biotec), and 0.1 IU/mL hIL21 (Miltenyi Biotec) in X-vivo 15 medium (Lonza) containing 2% Human Serum for 48h before being exposed twice to virus supernatant.

### Adoptive cell therapy

C57BL/6 HHD mice expressing the chimeric A2:K^b^ molecule (33) were bred at Erasmus MC and were used for adoptive T cell experiments as described previously with minor adaptations (24). Shortly, mice were transplanted with 0.5×10^6^ cells of the B16:HHD-YLE clone 18 to 25 days prior to T cell transfer at day 0. TCR T cells (6 to 7.5×10^6^ cells) were transferred intravenously, preceded by myeloablation through intraperitoneal injections of busulphan (16.5 µg/kg on days-4 and-3; Sigma) and cyclophosphamide (200 mg/kg on day-2; Sigma). For mock T cells, the same total number of cells was transferred as for wt TCR T cells. Tumor volumes were measured three times per week using a caliper and calculated using the formula 0.4 x (AxB^2^), where A represents the largest diameter measured, and B^2^ the perpendicular diameter (34). Tumor regression was defined as decreases in tumor volume of at least 30% after T cell transfer, whereas tumor relapse was defined as increases in tumor volume at 3 consecutive measurements after the initial tumor regression. Anti-tumor responses of mice at 115 days post-treatment were categorized as follows: a complete response as 100 % initial tumor regression with tumor remaining non-palpable; partial response as tumor regression followed by a tumor relapse; and non-response no tumor regression. 3-4 mice with regressing tumors 5 days after T cell transfer were sacrificed, and tumors were used for RNA sequencing (following snap freezing and storage at-80°C) as well as flow cytometric phenotyping of TILs (after making single cell suspensions). Mice were sacrificed when the tumor burden exceeded 1400 mm^3^, at which point relapsed tumors were collected. Blood samples were collected from the tail vein at weekly intervals for 11-16 weeks after T cell transfer and used for flow cytometric counting and phenotyping of TCR T cells.

### Detection of intracellular phosphorylation

Mouse T cells expressing either wt TCR or TCR:ICOS (6×10^4^ cells/well) were co-cultured with the B16:HHD-YLE clone or B16F10 wt (2×10^4^ cells/well) in Complete Mouse Medium in a tissue culture-treated 96-well plate at 37°C. After 2h, cells were fixed and stained and assessed for pPI3K (as described under ‘flow cytometry’). To assess the ability of TCR:ICOS to phosphorylate a large array of intracellular substrates, we treated wells from a non-tissue culture 96-well plate that were streptavidin pre-coated (Pierce) with 50µL of 1mg/mL biotinylated A2:Kb pentamer fused to the gp100 peptide YLEPGPVTA or an irrelevant peptide (FLYTYIAKV, Proimmune) for 1h at RT, after which wells were washed three times with PBS + 0.5% Tween-20 (Sigma). Mouse T cells expressing either gp100/A2 wt TCR, TCR:ICOS or TCR:ICOS-YF were added to the plates (0.25×10^6^ cells/well in 50µL Cytotox medium) for 30 min at 37°C, with frequencies of TCR-expressing T cells after transduction normalized towards TCR:ICOS T cells using Mock T cells. Following pMHC stimulations, plates were placed on ice and T cells were harvested and lysed with lysis beads, after which proteins were extracted, biotinylated and placed on the antibody array, followed by the addition of a Cy3-Streptavidin dye (GEPA43001, Merck) according to the manufacturer’s instructions (antibody array assay kit KAS02 and T cell Receptor Phospho-Antibody Array PTC188, Fullmoon Biosystems). To validate the effect of stimulation, one well from each condition was kept separate for flow cytometric detection of pERK1/2 (see above, and **Supplementary Figure 4b**). Detection of the Cy3 signal was performed using a Typhoon 5 fluorescent scanner and quantified using Imagequant TL with normalization according to the running ball background subtraction tool (Fiji, ImageJ 1.53t). Downstream analysis was performed using R, where median background of each slide was subtracted from the corresponding experimental values, and replicates (n=5) were averaged. Finally, the phosphorylation ratio after stimulation was calculated using the following formula:

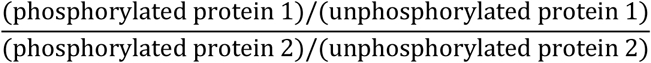

where 1 refers to the stimulated condition and 2 to the unstimulated condition. The different phosphorylated proteins were then classified into main T cell signaling pathways based on literature.

### Release of inflammatory cytokines

Mouse T cells expressing either wt TCR, TCR:ICOS or TCR:ICOS-YF (0.75×10^6^ cells/well) were co-cultured with the melanoma cell clone B16:HHD-YLE or B16F10 wt cells (0.25×10^6^ cells/well) in Complete Mouse Medium in a tissue culture treated 24-well plate for 16h at 37°C. Mock T cells served as negative controls. Supernatants were assessed for the presence of cytokines (according to a customized Luminex assay, R&D Systems Minneapolis USA; measuring IFN-γ, IL-1ß, IL-2, IL-4, IL-5, IL-6, IL-10, IL-17, CCL5, CCL20, CXCL2, MMP-8, MMP-9 and TNF-α). Values outside of range and not extrapolatable were excluded.

### Repeated stimulations with tumor cells

Mouse TCR T cells expressing either wt TCR, TCR:ICOS or TCR:ICOS-YF (0.75×10^6^) were co-cultured with B16:HHD-YLE or B16F10 wt cells (0.25×10^6^ cells) in Complete Mouse Medium supplemented with 25 IU/mL IL-2 in a 24-well plate for 16h at 37°C, with frequencies of TCR-expressing T cells after transduction normalized towards TCR:ICOS T cells using Mock T cells. After this 1st cycle of exposure, T cells were harvested with 20% of the sample being exposed to flow cytometric detection of T cell PD1 and TIM3, and 80% of the sample being again co-cultured for the 2nd cycle. Tumor cells were freshly pre-seeded to ensure an 80% confluency at start of co-culture with T cells. Experiments were performed over 4 cycles of 16h each. Repeated stimulation with human TCR T cells were performed in a similar manner, using either BLMgp100 (using gp100/A2 TCR T cells), MM-231-ROPN1 (ROPN1/A2 TCR) or MM-231-NY-ESO-1 (NY-ESO-1/A2 TCR) as target cells, freshly thawed T cells as effector cells, and co-culturing the cells in T cell medium supplemented with 50 IU/mL IL-2.

### Detection of antigen reactivity and sensitivity of TCR T cells

To assess antigen reactivity of murine TCR T cells, T cells expressing either wt TCR, or a co-stimulatory TCR (6×10^4^ cells/well) were co-cultured with B16-HHD-YLE in Cytotox medium in a tissue culture treated 96-well plate for 16h at 37°C. B16wt cells as well as Mock T cells were used as negative controls. To assess antigen sensitivity, TCR T cells (6×10^4^ cells/well) were co-cultured with B16-HHD cells (2×10^4^ cells/well) pretreated with 10µg/mL mIFN-γ (Sanquin) 24h prior to stimulation and loaded with cognate peptide (gp100 peptide) at increasing concentrations (from 0.1 nM to 30 µM) in Cytotox medium in a tissue culture treated 96-well plate for 16h at 37°C. Supernatants were assessed for the presence of mIFN-γ by enzyme-linked immunosorbent assay (ELISA, Biolegend). To assess antigen reactivity of human TCR T cells, T cells expressing either wt TCR or TCR:ICOS-RA (6×10^4^ cells/well) were co-cultured with BLM-gp100 (gp100/A2 TCR), MM-231-ROPN1 (ROPN1/A2 TCR) or MM-231-NY-ESO1 (NY-ESO1/A2 TCR) in Cytotox medium in a tissue culture treated 96-well plate for 16h at 37°C. BLM wt and MM-231 wt cell lines as well as Mock T cells were used as negative controls. To assess antigen sensitivity, TCR T cells (6×10^4^ cells/well) were co-cultured with T2 cells (2×10^4^ cells/well) loaded with cognate peptide (gp100, NY-ESO-1 or ROPN1 peptide) at increasing concentrations (from 0.1 nM to 30 µM) in Cytotox medium in a tissue culture treated 96-well plate for 16h at 37°C. Supernatants were assessed for the presence of hIFN-γ by enzyme-linked immunosorbent assay (ELISA, Biolegend).

### Detection of tumor cell killing ability of human TCR T cells

To assess killing ability of human TCR:ICOS T cells, BLM-gp100-eGFP (16×10^3^ cells), MM-231-ROPN1-eGFP (32×10^3^ cells) or MM-231-NY-ESO1-eGFP (16×10^3^ cells) were seeded onto wells from a flat bottom tissue culture treated 96 well plate (Falcon) in T cell medium supplemented with 50 IU/mL IL-2 overnight to reach a confluency of 50-80%. Human T cells expressing either wt TCR or hTCR:ICOS directed to either hgp100, NY-ESO1 or ROPN1 were thawed and co-cultured with their respective targets in an Incucyte SX5 placed inside a 37°C, 5% CO_2_ incubator (144×10^3^ cells/well targeting gp100, 96×10^3^ cells/well targeting ROPN1 and 32×10^3^ cells/well targeting NY-ESO1). Pictures of the wells were taken every 4h and GFP-covered areas were determined. Each experimental condition was run in quintuplicate and mock transduced T cells were used as negative controls.

### Statistical analyses

Statistical analysis and data display was performed using either GraphPad Prism (GraphPad Software, La Jolla, CA, version 8.0.2) or Rstudio (version 4.3.1). Statistical tests were considered significant when p<0.05. Changes in tumor volume or number of T cells over time were compared according to linear mixed models, with correction for random effect and elapsed time as a separate variable. All treatments were compared to the wt TCR condition and the interaction between treatment effect and time elapsed was reported. Survival of responding mice was compared according to the Cox regression model. Multiple group comparisons were performed either with 2-way ANOVA or, when the requirements were not met, Kruskal-Wallis test. Post-hoc testing was performed either with t-tests (if needed with Dunnets or Holm Sidak’s correction) or Dunns’ test.

For further description of materials and methods, particularly regarding cell culture, flow cytometry and RNA transcriptomics, see supplementary text.

## Results

### TCR:ICOS mediates durable anti-melanoma T cell responses

We designed a panel of co-stimulatory TCRs, all directed against the HLA-A2-restricted melanoma epitope gp100_280-288_. These new TCRα and β chains each consisted of the extracellular human variable and murine constant domains of the TCR, followed by the murine CD28 transmembrane domain, the intracellular domains of either murine CD28, CD134 (OX40), CD137 (4-1BB) or CD278 (ICOS) and the murine CD3ε intracellular domain (**Figure 1a**). This new design extended our previous observation that the transmembrane domain of CD28 and, downstream, the intracellular tail of CD3ε are beneficial for functional surface expression (24). All co-stimulatory TCRs were successfully transduced in murine T cells, except for the TCR with the 4-1BB domain, which showed very low surface expression, and was therefore excluded from further experiments (**Supplementary Figure 1a**). Co-stimulatory TCRs were next tested for antigen reactivity and sensitivity *in vitro*, which demonstrated antigen-specific IFN-γ production, although for all co-stimulatory TCR T cells with a lower avidity than wild-type (wt) TCR T cells (**Supplementary Figure 1b and c**). Subsequently, these TCRs were assessed in a mouse model for adoptive cell therapy, where B16:HHD-YLE melanoma cells overexpressing the cognate antigen were transplanted into immune competent HHD mice and T cells were transferred once the tumor became palpable (**Figure 1b**). Notably, while treatment with all TCR T cell products provided an initial tumor regression, only treatment with TCR:ICOS T cells provided an unambiguous advantage over wt TCR T cells (**Figure 1c**). In fact, treatment with TCR:ICOS T cells led to significantly increased survival, with 60% of the mice surviving as long as 150 days after T cell transfer compared to 0% for wt TCR and TCR:CD28 and 16% for TCR:OX40 T cells (**Figure 1d**). Moreover, mice treated with TCR:ICOS T cells showed a longer median time to relapse than those treated with other TCR T cells (140 days for TCR:ICOS vs 22, 32 and 35 days for wt TCR, TCR:OX40 and TCR:CD28, respectively, **Figure 1e**). Remarkably, when assessing the extent of response (i.e., complete, partial or non), TCR:ICOS T cell-treated mice had the highest rate of complete responders (i.e., 43% for TCR:ICOS vs 14, 0 and 0% for TCR:OX40, TCR:CD28 and wt TCR, **Supplementary Figure 1e)**. The durable anti-tumor response observed for mice treated with TCR:ICOS T cells was accompanied by a long-term persistence of circulating CD8 T cells expressing the introduced TCR (median of 119, 77 and 4 TCR+ CD8 T cells per µL blood at 100 days post-treatment for TCR:ICOS, TCR:OX40 and wt TCR, measurement not applicable for TCR:CD28 due to absence of surviving mouse **Figure 1f**). Of note, the relative long-term persistence observed with TCR:OX40 T cells was due to a single mouse showing persistent T cells. The enhanced persistence of TCR:ICOS T cells was also observed for CD4 T cells, albeit to a lesser extent (**Supplementary Figure 1d**). Taken together, these results demonstrate a superior *in vivo* durability of anti-melanoma responses, accompanied by a prolonged persistence of transferred T cells mediated by TCR:ICOS compared to alternative co-stimulatory TCRs.

### Intra-tumoral TCR:ICOS CD8 T cells exhibit a young and co-stimulatory phenotype

Next, we focused on TCR:ICOS and investigated whether its impact on the durability of anti-tumor responses was associated with the phenotype of circulating and tumor-infiltrating T cells, specifically their maturation state and expression of co-signaling receptors (schematically displayed in **Figure 2a**). When analyzing peripheral blood at 21 days post-treatment, we observed that TCR+ CD8 T cells from both TCR:ICOS and wt TCR T cell-treated mice were highly differentiated with the majority of cells expressing “antigen-experience” markers such as 4-1BB and displaying an effector memory phenotype with no clear differences between the two groups (**Figure 2b**). In addition, we observed low fractions of circulating TCR+ CD8 T cells co-expressing two or more co-inhibitory receptors and large fractions co-expressing two or more co-stimulatory receptors, again with no clear differences between the two treatment conditions. For instance, 16% of TCR:ICOS+ CD8 T cells and 17% of wt TCR+ CD8 T cells co-expressed PD1 and TIM3, whereas 76% of TCR:ICOS+ and 78% of wt TCR+ CD8 T cells co-expressed CD40L and 4-1BB. Also, circulating CD4 T cells showed similar patterns with little to no difference observed between treatment groups (**Supplementary Figure 2a**). In contrast, when analyzing regressing tumors at five days post-treatment, we observed that TCR:ICOS+ CD8 TILs exhibited a higher proportion of cells with a young naïve-like or central memory state and significantly fewer with a more differentiated effector memory state when compared to wt TCR+ CD8 TILs (**Figure 2c**, right panel). Notably, a larger proportion of TCR:ICOS+ CD8 TILs expressed 4-1BB and OX40, either as a single receptor, together or in combination with another co-stimulatory receptor relative to wt TCR+ CD8 TILs (**Figure 2c**, middle panel). With respect to CD4 TILs, treatment with TCR:ICOS T cells was characterized by a significantly higher fraction of cells expressing TIM3 and CTLA4 when compared to treatment with wt TCR+ T cells (**Supplementary Figure 2b)**. Moreover, we noted a higher frequency of co-stimulatory receptor-expressing TCR:ICOS+ CD4 TILs versus wt TCR+ CD4 TILs, even though differences did not reach statistical significance due to low numbers of TCR+ CD4 TILs measured. Collectively, these results show that while TCR:ICOS T cells are more abundant in the blood of mice, phenotypic differences of these cells were restricted to tumors, where TCR:ICOS+ CD8 TILs, but not wt TCR+ CD8 TILs, feature a young and co-stimulatory phenotype.

### TCR:ICOS T cells create an inflammatory micro-environment in tumors

Subsequently, we performed transcriptional profiling of tumors following treatment with either TCR:ICOS T cells or wt TCR T cells. Interestingly, genes that were differentially expressed in tumors that regressed after treatment with TCR:ICOS versus wt TCR T cells were to a large extent immune-related (e.g., Cxcl2, Il1b or Ccl5), indicative of a more inflammatory TME in TCR:ICOS-treated mice (**Figure 3a)**. In line with this, gene set enrichment analysis highlighted that “inflammatory response” was the most enriched pathway in regressing tumors from TCR:ICOS but not wt TCR T cell-treated mice (**Figure 3b)**. Additionally, the pathways “IL6 JAK STAT signaling” and “TNF-α signaling via NFκB” were also highly enriched in these tumors, further supporting the notion that TCR:ICOS T cells promote an inflammatory TME, at least in part through activation of the NFκB pathway. In contrast, transcriptomic analysis of relapsed tumors following treatment with TCR:ICOS T cells showed no significant enrichment of these or any other gene set compared to wt TCR T cells, indicating that treatment with TCR:ICOS T cells induced changes early-on, which were lost when tumors relapsed (**Supplementary Figure 3a**). In extension to this observation, the scores of various immune-related signatures that associate with either T cell infiltration, antigen presentation or T cell function (35,36) were reduced in relapsed versus regressed tumors (**Supplementary Figure 3b**). These findings, at the level of gene expression in tumors, suggest that TCR:ICOS T cells induce a local inflammatory TME.

### In vitro stimulated TCR:ICOS T cells activate the NFκB pathway, produce inflammatory cytokines and are resistant to exhaustion upon repeated tumor cell exposure

To test the mechanism of action of TCR:ICOS T cells toward the observed pro-inflammatory TME, we performed a series of *in vitro* assays. In preparation for these experiments, we confirmed activation-dependent phosphorylation of PI3K, a recognized target of ICOS, following stimulation with the tumor cell clone B16:HHD-YLE (the same as used for *in vivo* experiments, **Supplementary Figure 4a**). TCR:ICOS CD8 T cells showed significantly higher levels of phosphorylated PI3K (pPI3K) compared to wt TCR T cells. We also generated a TCR:ICOS-YF mutant in which the tyrosine of the active domain of ICOS was mutated into a phenylalanine, thereby abrogating activation of PI3K and its downstream substrates (**Figure 4a**) (25).

In a first series of experiments, T cells expressing either wt TCR, TCR:ICOS or TCR:ICOS-YF were stimulated with immobilized A2Kb chimeric MHC molecules coupled to the gp100 peptide or an irrelevant peptide for 30 minutes after which cell lysates were exposed to a kinase microarray. The occurrence of T cell activation was validated by detecting an increase in the fraction of TCR T cells positive for pERK1/2 (**Supplementary Figures 4b**). Remarkably, stimulation of TCR:ICOS T cells induced significant phosphorylation of multiple members of the NFκB family, such as Rel, p100/52, p105/50, IKKα/β, IKKγ and IKBβ, which was not observed in stimulated wt TCR or TCR:ICOS-YF T cells (**Figure 4b**). Expectedly, the PI3K pathway, with substrates such as AKT-Y236, is activated as well upon antigen-specific triggering of TCR:ICOS. The TCR or p38 pathways, harbor individual substrates that are also differentially phosphorylated when comparing TCR:ICOS to wt TCR T cells, however, when assessing pathways as a whole not to the same extent as the NFκB pathway. In a second series of experiments, we have stimulated TCR T cells with either antigen-positive or antigen-negative B16 tumor cells overnight and measured the secretion of 14 cytokines. We found that TCR:ICOS T cells produced significantly higher levels of cytokines and chemokines, such as IL-2, IL-6, CCL5, CXCL2 and TNF-α, in response to antigen-positive cells when compared to wt TCR T cells (**Figure 4c, Supplementary Figure 4c**). Importantly, this cytokine profile was not observed in T cells expressing the signaling-deficient TCR:ICOS-YF mutant. Many of the cytokines with enhanced secretion are direct downstream targets of the NFκB pathway and play active roles in tissue inflammation (37,38). Moreover, some of these mediators, such as IL-6, CCL5 and CXCL2, also showed increased gene expression in regressing tumors following treatment with TCR:ICOS T cells, demonstrating strong concordance between our *in vitro* and *in vivo* findings (**Figure 3a and b**). In a third series of experiments, we repeatedly exposed TCR T cells to fresh antigen-positive or negative melanoma cells and assessed the co-expression of PD1 and TIM3 as markers of T cell exhaustion. We found that the fraction of PD1^+^TIM3^+^ CD8 T cells over 3 rounds of antigen-specific stimulation was significantly lower for TCR:ICOS when compared to wt TCR or TCR:ICOS-YF T cells (**Figure 4d** and **Supplementary Figure 4d**). Taken together, this series of *in vitro* experiments revealed that TCR:ICOS preferentially signals through the PI3K-NFκB pathway, leading to the production of inflammatory cytokines and resulting in enhanced resistance to T cell exhaustion upon repeated stimulation. A proposed mechanism of action of TCR:ICOS is illustrated in **Supplementary Figure 4e**.

### Durability of anti-tumor response of T cells depends on ICOS-docking site for PI3K

To provide evidence that the PI3K pathway activation contributed to the *in vivo* efficacy of TCR:ICOS T cells, we compared the performance of TCR:ICOS-YF (the mutant unable to dock PI3K) to TCR:ICOS and wt TCR T cells in the melanoma mouse model (**Figure 5a**). Mice treated with TCR:ICOS T cells exhibited effective and durable antitumor response, whereas mice treated with either TCR:ICOS-YF or wt TCR T cells showed a substantially reduced survival (55, 30 and 20% survival at 150 days, respectively, **Figure 5b**, **5c** and **Supplementary Figure 5a**). In addition, median time to relapse was shorter in TCR:ICOS-YF and wt TCR T cells groups (30 and 26 days), while it was not reached within 150 days in the TCR:ICOS T cell group (**Figure 5d**). Lastly, TCR:ICOS CD8 T cells showed superior persistence in blood compared to TCR:ICOS-YF and wt TCR CD8 T cells (**Figure 5e**), whereas CD4 T cells exhibited lower persistence across all treatment groups (**Supplementary Figure 5b**). These results confirm that TCR:ICOS enhances the durability of anti-melanoma T cell responses through PI3K-dependent signaling.

### A single amino acid substitution in the cytosolic tail of the CD3ε domain enables functional expression of human TCR:ICOS in human T cells

In a final set of experiments, we translated our findings to a clinically relevant setting by designing and stringently testing a human TCR:ICOS construct (hTCR:ICOS) for its performance in human T cells. The rationale for these experiments comes from the observation that gene transfer of the gp100/A2 hTCR:ICOS using orthologous human instead of murine sequences for each building block resulted in low surface expression in human T cells (mean ± SEM: 5 ± 1% TCR-Vβ+ cells within CD3+ T cells, n=18) when compared to the murine TCR:ICOS construct (mTCR:ICOS, 17 ±3%, n=24) (**Supplementary Figure 6a and b**). To create a functionally expressed hTCR:ICOS while maintaining its overall format, we designed a panel of 50 variants to systemically test the impact of individual building blocks (or combinations thereof) and/or amino acid boundaries of these building blocks on surface expression in human T cells (**Figure 6a and Supplementary Table 1**). Using a stepwise approach, we started by exchanging each individual building block of the mTCR:ICOS (i.e., the extracellular TCR-C, the CD28 transmembrane, the ICOS intracellular and the CD3ε intracellular domains) for their human orthologues (**Figure 6b, left panel**). These mTCR:ICOS variants were introduced into human T cells, and surface expression was compared to the original mTCR:ICOS and wt TCR (**Figure 6b, middle panel**). While the introduction of human CD28 or ICOS domains did not impair expression, that of human TCR-C or CD3ε domains led to a notable lower expression (6 and 12% of TCR+ T cells, respectively, when compared to 18% for mTCR:ICOS). These results were re-assessed by performing the reverse experiment, where starting from hTCR:ICOS the latter two domains were substituted for their murine orthologues (**Figure 6b, right panel**). While introduction of the mTCR-C domain in an otherwise fully human hTCR:ICOS did not rescue the surface expression of hTCR:ICOS, introduction of the mCD3ε domain instead restored expression to a level comparable to that of mTCR:ICOS. In the next step, we focused on the five amino acid differences between human and murine CD3ε and constructed single amino acid substituents in hTCR:ICOS (**Figure 6c**). Strikingly, substitution of arginine at position 54 with alanine (hTCR:ICOS-CD3εR54A, in short: hTCR:ICOS-RA; corresponding to construct n°45 in **Supplementary Table 1**), led to expression levels that exceeded those of mTCR:ICOS and were similar to those of wt hTCR (35.5% for hTCR:ICOS-RA and wt hTCR vs 17.5% for mTCR:ICOS). In a last step, we validated the *in vitro* efficacy of this hTCR:ICOS-RA variant, characterizing its antigen sensitivity, reactivity, tumor killing ability as well as its resistance to T cell exhaustion. When challenged with titrated amounts of cognate peptide, hTCR:ICOS-RA T cells showed only slightly higher EC50 values and amplitude regarding IFN-γ production when compared to wt TCR T cells (**Figure 6d, 2^nd^ panel**). When stimulated with human melanoma cells that express the hgp100 antigen, hTCR:ICOS-RA T cells produced IFN-γ and effectively killed tumor cells (**Figure 6d**, **3^rd^ and 4^th^ panel**). Importantly, hTCR:ICOS-RA CD8 T cells showed significantly lower fractions co-expressing PD1 and TIM3 after repeated antigen stimulation compared to wt TCR T cells (**Figure 6d**, **5^th^ panel**).

Having established a successful format to gene-engineer human T cells with TCR:ICOS, we then evaluated whether this format could be extended to other TCR specificities targeting clinically relevant antigens. To this end, we tested surface expression and *in vitro* performance of hTCR:ICOS-RA directed against NY-ESO1/HLA-A2 and ROPN1/HLA-A2 (**Figure 6e** and **Figure 6f**). The NY-ESO1 TCR is directed against esophageal squamous cell carcinoma 1 antigen and is in an advanced stage of clinical development for synovial sarcoma (26), whereas the ROPN1 TCR targets Ropporin and is being developed for adoptive T cell therapy to treat Triple Negative Breast Cancer (TNBC) (27). For both NY-ESO1 and ROPN1 hTCR:ICOS, we demonstrated an R54A substitution-dependent surface expression (**Figure 6e and f, 1^st^ panels**). In addition, functional assays demonstrated that hTCR:ICOS-RA T cells have a similar antigen sensitivity relative to wt TCR T cells and robustly recognize and kill tumor cells. Lastly, and relevant to this study, hTCR:ICOS-RA CD8 T cells targeting either NY-ESO1 or ROPN1 also showed lower levels of T cell exhaustion markers (PD1 and TIM3) following multiple rounds of repeated antigen stimulation when compared to their wt TCR counterparts. In short, we found that surface expression of hTCR:ICOS was dependent on a single amino acid substitution, which preserved antigen sensitivity and recognition as well as resistance to the induction of T cell exhaustion of TCRs with multiple antigen specificities.

## Discussion

Our current study unveils a new TCR format that provides built-in co-stimulation through ICOS and enables exceptionally prolonged anti-tumor T cell responses in an immune competent melanoma mouse model. TCR:ICOS T cells show a young and co-stimulatory cell state and are responsible for an inflammatory TME. These characteristics are recapitulated *in vitro*, where these T cells signal through the PI3K and NFκB pathways, produce inflammatory cytokines and do not become exhausted upon repeated tumor cell exposure. These advantages were shown to be reversed when disrupting ICOS-mediated activation of PI3K. Surface expression of TCR:ICOS in human T cells strictly requires a specific single amino acid substitution in the intracellular tail, which does not limit antigen sensitivity nor reactivity, yet preserves the TCR’s ability to limit expression of the immune checkpoints PD1 and TIM3. Notably, the optimized human ‘ICOS cassette’ was able to sustain the anti-tumor performance of human T cells across multiple antigen specificities TCRs that are genetically coupled to co-stimulatory molecules are conceptually similar to CARs, where incorporation of a co-stimulatory molecule is standard practice. However, engineering of TCRs and CARs differs significantly. In fact, TCRs face more stringent structural constraints due to the complex and step-wise assembly of TCRα and TCRß chains with CD3 components as well as the requirement for precise binding to peptide:MHC (39–41). Consequently, most studies so far have relied on alternative concepts, such as TCRs that are supported by a second chimeric co-stimulatory receptor (42), where improved T cell function depends on two separate receptors. We designed and tested mouse TCRs with built-in intracellular domains of either CD28, OX40 or ICOS, using, as a starting point, a chimeric TCR format with the transmembrane domain of CD28 and the cytosolic domain of CD3ε (24). A TCR incorporating the 4-1BB molecule did not show surface expression on mouse T cells, possibly due to the tendency of murine 4-1BB to be expressed as a monomer or due to its basic amino acid-rich intracellular motif that was shown to be detrimental to surface expression in other settings (43–45). In fact, 4-1BB has been incorporated successfully into CARs and chimeric co-receptors, but, as far we are aware, not into TCRs (42,46,47). When mice were treated with TCR:OX40, TCR:CD28 and wt TCR T cells, they showed initial anti-tumor responses; however, relapse of tumors ensued within 20 to 35 days, mainly resulting in no or partial responses at the end of the experiments. In contrast, when mice were treated with TCR:ICOS T cells, tumor relapses were delayed up to 140 days and over 40% of mice showed complete responses. This anti-tumor response was accompanied by a significantly enhanced persistence of TCR:ICOS-expressing CD8 T cells in the blood of mice. These findings are concordant with those reported for CAR:ICOS T cells (48–51), where these T cells also showed long-term persistence and a durable anti-tumor effect. Interestingly, although circulating TCR:ICOS T cells were more abundant than wt TCR T cells, no difference was found with respect to their phenotype. Within tumors, however, CD8 TILs that expressed TCR:ICOS exhibited a less differentiated and more co-stimulatory phenotype than CD8 TILs that expressed wt TCR, with close to half of the former cells harboring a naïve-like or central memory state and significantly more cells expressing 4-1BB and OX40. This young, yet most likely antigen-experienced and activated phenotype of TCR:ICOS T cells likely contributes to the improved T cell persistence and anti-tumor response we observed in our mouse model, consistent with prior reports linking less differentiated T cells to improved long-term functionality (52–55). Our current flow cytometry analysis did not distinguish between naïve-like and stem cell-like T cell subsets, which warrants further investigation (56). Besides, recent studies have also highlighted the critical role of ICOS signaling in the formation of CD8 resident memory T cells (CD8 Trm), which are recognized key players for long-term protection against tumor relapse (57–59). Although we did not specifically stain for Trm markers, it is unlikely that these cells were present at the time of tumor isolation since Peng and colleagues have shown that CD8 Trm cells are not present yet five days after T cell transfer, and only begin to emerge around days 7-8, peaking no earlier than 40 days after adoptive transfer (57). Future studies could assess whether TCR:ICOS promotes Trm differentiation and investigate the kinetics of these cells.

Treatment with TCR:ICOS T cells initiated a significant enrichment of inflammation-related gene sets in regressing tumors, particularly those related to the NFκB pathway. This was recapitulated *in vitro,* where antigen-stimulated TCR:ICOS T cells, but not wt TCR T cells, phosphorylated multiple components of the NFκB pathway and produced significantly higher levels of inflammatory cytokines that are dependent upon NFκB activation, such as CCL5, IL-2, IL-6 and TNF-α (60–64). These effects were abolished in T cells expressing TCR:ICOS-YF, a mutant of ICOS that does not dock nor activate PI3K, proving that activation of the NFκB pathway and production of inflammatory cytokines is PI3K-dependent. Our observations, particularly the ICOS-mediated increase in TNF-α secretion, extend previous reports (65,66). Moreover, TNF-α was also shown to up-regulate expression of ICOS-L by antigen presenting cells, thereby inducing a positive feedback loop, which may further contribute to the durable responses of TCR:ICOS T cells (67). In contrast, associations between ICOS stimulation and IL-2, IL-6, CCL5 and CXCL2 secretion were previously less well established (68–74). Interestingly, CAR:ICOS also mediates increased antigen-specific production of TNF-α, IL-2 and IFN-γ compared to first-generation CARs that lack co-stimulatory molecules (49,50). CAR:ICOS mediates production of IL-17 as well, a feature not observe with TCR:ICOS. This discrepancy is somewhat unexpected given that both ICOS and NFκB activation have previously been linked to Th17 differentiation (37,75–78). One explanation may be that CD4 T cell enrichments have been performed for the generation of CAR:ICOS T cells, but not TCR:ICOS T cells. An alternative explanation may relate to differences in receptor affinity and/or signaling since CARs bind antigen with high affinity whereas TCRs bind antigen with moderate affinity and CARs are coupled to CD3ζ whereas TCRs in this study are coupled to CD3ε, potentially leading to differential activation of molecules such as LAT and Lck. In addition to the activation of NFκB and production of inflammatory cytokines, we also observed that TCR:ICOS provided resistance to PD1 and TIM3 co-expression by T cells upon repeated stimulation with antigen-positive tumor cells, again depending on PI3K. Importantly, a similar observation was also made in our mouse model, where we found that mice treated with TCR:ICOS-YF T cells did not demonstrate durable anti-tumor responses, underscoring the critical role of PI3K-NFκB signaling in mediating the beneficial effects of TCR:ICOS T cells. It is noteworthy that while CD28 also activates the PI3K and NFκB pathways, ICOS is reported to more strongly recruit the p50α regulatory component of PI3K, resulting in greater downstream Akt phosphorylation (79). This would indicate that the superior performance of TCR:ICOS may stem from a specific sequence involving PI3K and Akt activation and culminating in the engagement of NFκB (summarized in **Supplementary Figure 4e**).

Following determination of the therapeutic value of TCR:ICOS and identification of its mechanism of action, we set out to translate TCR:ICOS to human T cells. Murine TCR:ICOS (mTCR:ICOS) was expressed on the surface of human T cells, albeit at a low level, yet a fully murine sequence is clinically undesirable due to its risk of inducing xenoreactivity. However, the fully humanized orthologue of TCR:ICOS (hTCR:ICOS) was not expressed on the surface of human T cells. Through systematic testing of 50 constructs, we identified a single amino acid - arginine at position 54 in the intracellular tail of hCD3ε - as the key barrier to surface expression of hTCR:ICOS. Replacing this residue with an alanine (as found in the murine counterpart) completely rescued surface expression of hTCR:ICOS on human T cells. One possible explanation is that the positively charged arginine in the tail of hCD3ε acts as a retention signal, preventing hTCR:ICOS from exiting the endoplasmic reticulum (ER) and reaching the plasma membrane. In fact, previous studies have reported that this amino acid plays an integral role in an ER retention signal to guide and facilitate correct assembly of TCR/CD3 components in T cells (80,81). The identification of a surface-expressed hTCR:ICOS-RA variant was critical in translating this new chimeric TCR toward clinical application. To expand the bandwidth of this new TCR, we also applied the optimized hTCR:ICOS format to TCRs with other antigen specificities, namely NY-ESO1 and ROPN-1. Once more, the R54A substitution proved both necessary and sufficient to restore surface expression of hTCR:ICOS. Functional testing across all three specificities demonstrated that the optimized hTCR:ICOS did not compromise antigen sensitivity nor T cell effector function. This was exemplified through peptide titration assays, as well as tumor cell-specific IFN-γ production and cytotoxicity, all of which were comparable to wt TCR T cells. These conventional *in vitro* T cell assays have traditionally been used to guide selection of TCRs for clinical development (82). However, these assays primarily measure immediate effector function and may not reflect the long-term durability of T cell responses. The repeated stimulation assay may serve as a valuable complementary *in vitro* tool for evaluating the sustained functionality of engineered human TCRs. Importantly, the optimized hTCR:ICOS did confer a clear and significant advantage in terms of resisting T cell exhaustion in this assay across all TCR specificities.

## Conclusions

In conclusion, we introduce a TCR format with built-in ICOS that enhances T cell fitness upon exposure to antigen. This TCR:ICOS triggers the PI3K-NFκB pathway, drives an inflammatory tumor milieu, reduces T cell exhaustion, and ultimately leads to enhanced T cell persistence and long-term tumor clearance. In addition, we have developed an optimized variant of TCR:ICOS that supports robust functional expression in human T cells while preserving its therapeutic benefits across multiple TCR specificities. Taken together, the TCR:ICOS format may constitute a promising platform for adoptive T cell therapy. In fact, preparations are currently underway for a first-in-human phase 1b trial evaluating TCR:ICOS T cells targeting the ROPN-1 antigen in patients with TNBC, marking a pivotal step toward clinical translation and paving the way for next-generation TCR-based immunotherapies.

## Declarations

### Ethics approval

Mice experiments were approved by the Experimental Animal Committee of the Erasmus MC Cancer Institute and carried out in accordance with institutional and national guidelines.

## Consent for publication

Not applicable.

## Availability of data and materials

The datasets used during the current study are available from the corresponding author on reasonable request.

## Authors’ Contributions

**AM:** Conceptualization, data curation, formal analysis, investigation, visualization, methodology, writing and editing. **CB:** Conceptualization, data curation, investigation, methodology. **MM:** Investigation, data curation, methodology. **RW**: Investigation. **DR:** Investigation. **MvB:** Investigation. **KK:** Investigation. **M.J.W.P:** Investigation, data curation, methodology. **W.A.D:** Investigation. **AK:** data curation, investigation, methodology. **DH:** Conceptualization, writing and editing. **R.J.M.A:** funding acquisition, writing and editing. **CS:** Conceptualization, methodology, supervision, writing and editing. **RD:** Conceptualization, methodology, resources, supervision, funding acquisition, writing and editing. All authors read and approved the final manuscript.

## Acknowledgements

The authors thank Coen Govers, Rachid Bouzid, Luc Veeneman and Ana Avramovic, who participated in this work. Furthermore, we would like to thank the many healthy blood donors from Sanquin whose donation allowed the translations of our findings to the human setting.

## List of abbreviations

CAR: chimeric antigen receptor
CD: cluster of differentiation
CCL: Chemokine (C-C motif) ligand
CMM: Complete mouse medium
CXCL: Chemokine (C-X-C motif)
DNA: Deoxyribonucleic acid
EC: Extracellular
ER: Endoplasmic reticulum
ELISA: Enzyme linked immunosorbent assay
FDA: Food and drug administration
GFP: Green fluorescent protein
HLA: Human leukocyte antigen
IC: Intracellular
ICOS: Inducible T cell co-stimulator
IFN-γ: Interferon gamma
IL: Interleukin
IKK: IkappaB-kinase
MHC: Major histocompatibility complex
MMP: matrix metalloproteinase
NF-κB: Nuclear factor kappa B
NY-ESO1: New-York esophageal squamous cell carcinoma 1
PBMC: peripheral blood mononuclear cell
PCR: Polymerase chain reaction
PD1: programmed cell death protein 1
PI3K: Phosphoinositide 3-kinase
RNA: Ribonucleic acid
ROPN1: Rhophilin associated tail protein 1
RT: Room temperature
TCR: T cell receptor
Th17: T helper 17
TIL: Tumor infiltrating lymphocyte
TIM3: T cell immunoglobulin mucin-3
TM: Transmembrane
TME: Tumor micro-environment
TNBC: Triple negative breast cancer
TNF-α: Tumor necrosis factor alpha
Trm: Tissue resident memory (T cell)
wt: Wildtype

## References

1. Kochenderfer JN, Dudley ME, Carpenter RO, Kassim SH, Rose JJ, Telford WG, et al. Donor-derived CD19-targeted T cells cause regression of malignancy persisting after allogeneic hematopoietic stem cell transplantation. Blood. 2013 Dec 12;122(25):4129–39.

2. Brentjens RJ, Davila ML, Riviere I, Park J, Wang X, Cowell LG, et al. CD19-targeted T cells rapidly induce molecular remissions in adults with chemotherapy-refractory acute lymphoblastic leukemia. Sci Transl Med. 2013 Mar 20;5(177):177ra38.

3. Schuster SJ, Bishop MR, Tam CS, Waller EK, Borchmann P, McGuirk JP, et al. Tisagenlecleucel in Adult Relapsed or Refractory Diffuse Large B-Cell Lymphoma. N Engl J Med. 2019 Jan 3;380(1):45–56.

4. Porter DL, Levine BL, Kalos M, Bagg A, June CH. Chimeric Antigen Receptor–Modified T Cells in Chronic Lymphoid Leukemia. New England Journal of Medicine. 2011 Aug 25;365(8):725–33.

5. Neelapu SS, Locke FL, Bartlett NL, Lekakis LJ, Miklos DB, Jacobson CA, et al. Axicabtagene Ciloleucel CAR T-Cell Therapy in Refractory Large B-Cell Lymphoma. New England Journal of Medicine. 2017 Dec 28;377(26):2531–44.

6. Locke FL, Ghobadi A, Jacobson CA, Miklos DB, Lekakis LJ, Oluwole OO, et al. Long-term safety and activity of axicabtagene ciloleucel in refractory large B-cell lymphoma (ZUMA-1): a single-arm, multicentre, phase 1–2 trial. The Lancet Oncology. 2019 Jan 1;20(1):31–42.

7. Moretti A, Ponzo M, Nicolette CA, Tcherepanova IY, Biondi A, Magnani CF. The Past, Present, and Future of Non-Viral CAR T Cells. Front Immunol. 2022;13:867013.

8. Sarnaik AA, Hamid O, Khushalani NI, Lewis KD, Medina T, Kluger HM, et al. Lifileucel, a Tumor-Infiltrating Lymphocyte Therapy, in Metastatic Melanoma. J Clin Oncol. 2021 Aug 20;39(24):2656–66.

9. Hong DS, Van Tine BA, Biswas S, McAlpine C, Johnson ML, Olszanski AJ, et al. Autologous T cell therapy for MAGE-A4+ solid cancers in HLA-A*02+ patients: a phase 1 trial. Nat Med. 2023 Jan;29(1):104–14.

10. Kishton RJ, Restifo NP. T cells lead the charge against solid tumors. Nat Cancer. 2024 Dec;5(12):1762–4.

11. Debets R, Donnadieu E, Chouaib S, Coukos G. TCR-engineered T cells to treat tumors: Seeing but not touching? Semin Immunol. 2016 Feb;28(1):10–21.

12. Hammerl D, Rieder D, Martens JWM, Trajanoski Z, Debets R. Adoptive T Cell Therapy: New Avenues Leading to Safe Targets and Powerful Allies. Trends in Immunology. 2018 Nov 1;39(11):921–36.

13. Chandran SS, Klebanoff CA. T cell receptor-based cancer immunotherapy: Emerging efficacy and pathways of resistance. Immunol Rev. 2019 Jul;290(1):127–47.

14. Capece D, Verzella D, Fischietti M, Zazzeroni F, Alesse E. Targeting Costimulatory Molecules to Improve Antitumor Immunity. BioMed Research International. 2012;2012(1):926321.

15. Franzese O. Tumor Microenvironment Drives the Cross-Talk Between Co-Stimulatory and Inhibitory Molecules in Tumor-Infiltrating Lymphocytes: Implications for Optimizing Immunotherapy Outcomes. International Journal of Molecular Sciences. 2024 Jan;25(23):12848.

16. Spranger S, Spaapen RM, Zha Y, Williams J, Meng Y, Ha TT, et al. Up-Regulation of PD-L1, IDO, and Tregs in the Melanoma Tumor Microenvironment Is Driven by CD8+ T Cells. Science Translational Medicine. 2013 Aug 28;5(200):200ra116-200ra116.

17. Gyurdieva A, Zajic S, Chang YF, Houseman EA, Zhong S, Kim J, et al. Biomarker correlates with response to NY-ESO-1 TCR T cells in patients with synovial sarcoma. Nat Commun. 2022 Sep 8;13(1):5296.

18. Galon J, Costes A, Sanchez-Cabo F, Kirilovsky A, Mlecnik B, Lagorce-Pagès C, et al. Type, Density, and Location of Immune Cells Within Human Colorectal Tumors Predict Clinical Outcome. Science. 2006 Sep 29;313(5795):1960–4.

19. Hammerl D, Martens JWM, Timmermans M, Smid M, Trapman-Jansen AM, Foekens R, et al. Spatial immunophenotypes predict response to anti-PD1 treatment and capture distinct paths of T cell evasion in triple negative breast cancer. Nat Commun. 2021 Sep 27;12(1):5668.

20. Schaft N, Willemsen RA, de Vries J, Lankiewicz B, Essers BWL, Gratama JW, et al. Peptide fine specificity of anti-glycoprotein 100 CTL is preserved following transfer of engineered TCR alpha beta genes into primary human T lymphocytes. J Immunol. 2003 Feb 15;170(4):2186–94.

21. Pouw NMC, Westerlaken EJ, Willemsen RA, Debets R. Gene transfer of human TCR in primary murine T cells is improved by pseudo-typing with amphotropic and ecotropic envelopes. J Gene Med. 2007 Jul;9(7):561–70.

22. Engels B, Cam H, Schüler T, Indraccolo S, Gladow M, Baum C, et al. Retroviral vectors for high-level transgene expression in T lymphocytes. Hum Gene Ther. 2003 Aug 10;14(12):1155–68.

23. Straetemans T, Berrevoets C, Coccoris M, Treffers-Westerlaken E, Wijers R, Cole DK, et al. Recurrence of Melanoma Following T Cell Treatment: Continued Antigen Expression in a Tumor That Evades T Cell Recruitment. Mol Ther. 2015 Feb;23(2):396–406.

24. Govers C, Sebestyén Z, Roszik J, van Brakel M, Berrevoets C, Szöőr Á, et al. TCRs genetically linked to CD28 and CD3ε do not mispair with endogenous TCR chains and mediate enhanced T cell persistence and anti-melanoma activity. J Immunol. 2014 Nov 15;193(10):5315–26.

25. Gigoux M, Shang J, Pak Y, Xu M, Choe J, Mak TW, et al. Inducible costimulator promotes helper T-cell differentiation through phosphoinositide 3-kinase. Proc Natl Acad Sci U S A. 2009 Dec 1;106(48):20371–6.

26. D’Angelo SP, Melchiori L, Merchant MS, Bernstein D, Glod J, Kaplan R, et al. Antitumor Activity Associated with Prolonged Persistence of Adoptively Transferred NY-ESO-1 c259T Cells in Synovial Sarcoma. Cancer Discov. 2018 Aug;8(8):944–57.

27. Kortleve D, Hammerl D, van Brakel M, Wijers R, Roelofs D, Kroese K, et al. TCR-engineered T-cells directed against Ropporin-1 constitute a safe and effective treatment for triple-negative breast cancer. Cancer Discov. 2024 Aug 22;

28. Li Y, Moysey R, Molloy PE, Vuidepot AL, Mahon T, Baston E, et al. Directed evolution of human T-cell receptors with picomolar affinities by phage display. Nat Biotechnol. 2005 Mar;23(3):349–54.

29. Soneoka Y, Cannon PM, Ramsdale EE, Griffiths JC, Romano G, Kingsman SM, et al. A transient three-plasmid expression system for the production of high titer retroviral vectors. Nucleic Acids Res. 1995 Feb 25;23(4):628–33.

30. Lamers CHJ, Willemsen RA, Luider BA, Debets R, Bolhuis RLH. Protocol for gene transduction and expansion of human T lymphocytes for clinical immunogene therapy of cancer. Cancer Gene Ther. 2002 Jul;9(7):613–23.

31. Weijtens ME, Willemsen RA, Hart EH, Bolhuis RL. A retroviral vector system ‘STITCH’ in combination with an optimized single chain antibody chimeric receptor gene structure allows efficient gene transduction and expression in human T lymphocytes. Gene Ther. 1998 Sep;5(9):1195–203.

32. Lamers CHJ, van Steenbergen-Langeveld S, van Brakel M, Groot-van Ruijven CM, van Elzakker PMML, van Krimpen B, et al. T Cell Receptor-Engineered T Cells to Treat Solid Tumors: T Cell Processing Toward Optimal T Cell Fitness. Hum Gene Ther Methods. 2014 Dec 1;25(6):345–57.

33. Pascolo S, Bervas N, Ure JM, Smith AG, Lemonnier FA, Pérarnau B. HLA-A2.1-restricted education and cytolytic activity of CD8(+) T lymphocytes from beta2 microglobulin (beta2m) HLA-A2.1 monochain transgenic H-2Db beta2m double knockout mice. J Exp Med. 1997 Jun 16;185(12):2043–51.

34. Seynhaeve ALB, Hoving S, Schipper D, Vermeulen CE, de Wiel-Ambagtsheer G aan, van Tiel ST, et al. Tumor necrosis factor alpha mediates homogeneous distribution of liposomes in murine melanoma that contributes to a better tumor response. Cancer Res. 2007 Oct 1;67(19):9455–62.

35. Hammerl D, Massink MPG, Smid M, van Deurzen CHM, Meijers-Heijboer HEJ, Waisfisz Q, et al. Clonality, Antigen Recognition, and Suppression of CD8+ T Cells Differentially Affect Prognosis of Breast Cancer Subtypes. Clin Cancer Res. 2020 Jan 15;26(2):505–17.

36. Kortleve D, Coelho RML, Hammerl D, Debets R. Cancer germline antigens and tumor-agnostic CD8+ T cell evasion. Trends Immunol. 2022 May;43(5):391–403.

37. Liu T, Zhang L, Joo D, Sun SC. NF-κB signaling in inflammation. Signal Transduct Target Ther. 2017 Jul 14;2:17023.

38. Su CM, Wang L, Yoo D. Activation of NF-κB and induction of proinflammatory cytokine expressions mediated by ORF7a protein of SARS-CoV-2. Sci Rep. 2021 Jun 29;11(1):13464.

39. Geisler C, Kuhlmann J, Rubin B. Assembly, intracellular processing, and expression at the cell surface of the human alpha beta T cell receptor/CD3 complex. Function of the CD3-zeta chain. Journal of Immunology. 1989;143(12):4069–77.

40. Kappes DJ, Tonegawa S. Surface expression of alternative forms of the TCR/CD3 complex. Proceedings of the National Academy of Sciences. 1991 Dec;88(23):10619–23.

41. Hall C, Berkhout B, Alarcon B, Sancho J, Wileman T, Terhorst C. Requirements for cell surface expression of the human TCR/CD3 complex in non-T cells. Int Immunol. 1991 Apr;3(4):359–68.

42. Hernández-López P, van Diest E, Brazda P, Heijhuurs S, Meringa A, Hoorens van Heyningen L, et al. Dual targeting of cancer metabolome and stress antigens affects transcriptomic heterogeneity and efficacy of engineered T cells. Nat Immunol. 2024 Jan;25(1):88–101.

43. Lah S, Kim S, Kang I, Kim H, Hupperetz C, Jung H, et al. Engineering second-generation TCR-T cells by site-specific integration of TRAF-binding motifs into the CD247 locus. J Immunother Cancer. 2023 Apr 1;11(4):e005519.

44. Chin SM, Kimberlin CR, Roe-Zurz Z, Zhang P, Xu A, Liao-Chan S, et al. Structure of the 4-1BB/4-1BBL complex and distinct binding and functional properties of utomilumab and urelumab. Nat Commun. 2018 Nov 8;9(1):4679.

45. Bitra A, Doukov T, Destito G, Croft M, Zajonc DM. Crystal structure of the m4-1BB/4-1BBL complex reveals an unusual dimeric ligand that undergoes structural changes upon 4-1BB receptor binding. J Biol Chem. 2019 Feb 8;294(6):1831–45.

46. Milone MC, Fish JD, Carpenito C, Carroll RG, Binder GK, Teachey D, et al. Chimeric receptors containing CD137 signal transduction domains mediate enhanced survival of T cells and increased antileukemic efficacy in vivo. Mol Ther. 2009 Aug;17(8):1453–64.

47. Zhi L, Yin M, Su X, Zhang Z, Lu H, Li M, et al. A chimeric switch-receptor PD1-DAP10-41BB augments NK92-cell activation and killing for human lung Cancer H1299 Cell. Biochemical and Biophysical Research Communications. 2022 Apr 16;600:94–100.

48. Shen CJ, Yang YX, Han EQ, Cao N, Wang YF, Wang Y, et al. Chimeric antigen receptor containing ICOS signaling domain mediates specific and efficient antitumor effect of T cells against EGFRvIII expressing glioma. Journal of Hematology & Oncology. 2013 May 9;6(1):33.

49. Guedan S, Chen X, Madar A, Carpenito C, McGettigan SE, Frigault MJ, et al. ICOS-based chimeric antigen receptors program bipolar TH17/TH1 cells. Blood. 2014 Aug 14;124(7):1070–80.

50. Guedan S, Posey AD, Shaw C, Wing A, Da T, Patel PR, et al. Enhancing CAR T cell persistence through ICOS and 4-1BB costimulation. JCI Insight. 2018 Jan 11;3(1):e96976, 96976.

51. Guedan S, Madar A, Casado-Medrano V, Shaw C, Wing A, Liu F, et al. Single residue in CD28-costimulated CAR-T cells limits long-term persistence and antitumor durability. 2020 Jun 1;

52. Tantalo DG, Oliver AJ, von Scheidt B, Harrison AJ, Mueller SN, Kershaw MH, et al. Understanding T cell phenotype for the design of effective chimeric antigen receptor T cell therapies. J Immunother Cancer. 2021 May 25;9(5):e002555.

53. Klebanoff CA, Gattinoni L, Palmer DC, Muranski P, Ji Y, Hinrichs CS, et al. Determinants of Successful CD8+ T-Cell Adoptive Immunotherapy for Large Established Tumors in Mice. Clinical Cancer Research. 2011 Aug 14;17(16):5343–52.

54. Kwon BS, Weissman SM. cDNA sequences of two inducible T-cell genes. Proc Natl Acad Sci U S A. 1989 Mar;86(6):1963–7.

55. Fujita T, Ukyo N, Hori T, Uchiyama T. Functional characterization of OX40 expressed on human CD8+ T cells. Immunology Letters. 2006 Jul 15;106(1):27–33.

56. Gattinoni L, Klebanoff CA, Restifo NP. Paths to stemness: building the ultimate antitumour T cell. Nat Rev Cancer. 2012 Oct;12(10):671–84.

57. Peng C, Huggins MA, Wanhainen KM, Knutson TP, Lu H, Georgiev H, et al. Engagement of the costimulatory molecule ICOS in tissues promotes establishment of CD8+ tissue-resident memory T cells. Immunity. 2022 Jan 11;55(1):98–114.e5.

58. Damei I, Trickovic T, Mami-Chouaib F, Corgnac S. Tumor-resident memory T cells as a biomarker of the response to cancer immunotherapy. Front Immunol. 2023 Jul 20;14.

59. Okła K, Farber DL, Zou W. Tissue-resident memory T cells in tumor immunity and immunotherapy. J Exp Med. 2021 Apr 5;218(4):e20201605.

60. Takeuchi O, Akira S. Pattern Recognition Receptors and Inflammation. Cell. 2010 Mar 19;140(6):805–20.

61. Wickremasinghe MI, Thomas LH, O’Kane CM, Uddin J, Friedland JS. Transcriptional mechanisms regulating alveolar epithelial cell-specific CCL5 secretion in pulmonary tuberculosis. J Biol Chem. 2004 Jun 25;279(26):27199–210.

62. Hoyos B, Ballard DW, Böhnlein E, Siekevitz M, Greene WC. Kappa B-specific DNA binding proteins: role in the regulation of human interleukin-2 gene expression. Science. 1989 Apr 28;244(4903):457–60.

63. Libermann TA, Baltimore D. Activation of interleukin-6 gene expression through the NF-kappa B transcription factor. Mol Cell Biol. 1990 May;10(5):2327–34.

64. Collart MA, Baeuerle P, Vassalli P. Regulation of tumor necrosis factor alpha transcription in macrophages: involvement of four kappa B-like motifs and of constitutive and inducible forms of NF-kappa B. Mol Cell Biol. 1990 Apr;10(4):1498–506.

65. Li J, Heinrichs J, Leconte J, Haarberg K, Semple K, Liu C, et al. Phosphatidylinositol 3-Kinase–Independent Signaling Pathways Contribute to ICOS-Mediated T Cell Costimulation in Acute Graft-Versus-Host Disease in Mice. J Immunol. 2013 Jul 1;191(1):200–7.

66. Sainson RCA, Thotakura AK, Kosmac M, Borhis G, Parveen N, Kimber R, et al. An Antibody Targeting ICOS Increases Intratumoral Cytotoxic to Regulatory T-cell Ratio and Induces Tumor Regression. Cancer Immunol Res. 2020 Dec;8(12):1568–82.

67. Richter G, Hayden-Ledbetter M, Irgang M, Ledbetter JA, Westermann J, Körner I, et al. Tumor Necrosis Factor-α Regulates the Expression of Inducible Costimulator Receptor Ligand on CD34+ Progenitor Cells during Differentiation into Antigen Presenting Cells*. Journal of Biological Chemistry. 2001 Dec 7;276(49):45686–93.

68. Odegard JM, DiPlacido LD, Greenwald L, Kashgarian M, Kono DH, Dong C, et al. ICOS Controls Effector Function but Not Trafficking Receptor Expression of Kidney-Infiltrating Effector T Cells in Murine Lupus. J Immunol. 2009 Apr 1;182(7):4076–84.

69. Alves GF, Stoppa I, Aimaretti E, Monge C, Mastrocola R, Porchietto E, et al. ICOS-Fc as innovative immunomodulatory approach to counteract inflammation and organ injury in sepsis. Front Immunol. 2022 Sep 2;13.

70. Riley JL, Blair PJ, Musser JT, Abe R, Tezuka K, Tsuji T, et al. ICOS Costimulation Requires IL-2 and Can Be Prevented by CTLA-4 Engagement1. The Journal of Immunology. 2001 Apr 15;166(8):4943–8.

71. Harada Y, Ohgai D, Watanabe R, Okano K, Koiwai O, Tanabe K, et al. A single amino acid alteration in cytoplasmic domain determines IL-2 promoter activation by ligation of CD28 but not inducible costimulator (ICOS). J Exp Med. 2003 Jan 20;197(2):257–62.

72. Hutloff A, Dittrich AM, Beier KC, Eljaschewitsch B, Kraft R, Anagnostopoulos I, et al. ICOS is an inducible T-cell co-stimulator structurally and functionally related to CD28. Nature. 1999 Dec;402(6763):21–4.

73. Yoshinaga SK, Zhang M, Pistillo J, Horan T, Khare SD, Miner K, et al. Characterization of a new human B7-related protein: B7RP-1 is the ligand to the co-stimulatory protein ICOS. Int Immunol. 2000 Oct;12(10):1439–47.

74. Yoshinaga SK, Whoriskey JS, Khare SD, Sarmiento U, Guo J, Horan T, et al. T-cell co-stimulation through B7RP-1 and ICOS. Nature. 1999 Dec;402(6763):827–32.

75. Park SH, Cho G, Park SG. NF-κB Activation in T Helper 17 Cell Differentiation. Immune Netw. 2014 Feb;14(1):14–20.

76. Molinero LL, Cubre A, Mora-Solano C, Wang Y, Alegre ML. T cell receptor/CARMA1/NF-κB signaling controls T-helper (Th) 17 differentiation. Proceedings of the National Academy of Sciences. 2012 Nov 6;109(45):18529–34.

77. Wyatt MM, Huff LW, Nelson MH, Neal LR, Medvec AR, Rangel Rivera GO, et al. Augmenting TCR signal strength and ICOS costimulation results in metabolically fit and therapeutically potent human CAR Th17 cells. Mol Ther. 2023 Jul 5;31(7):2120–31.

78. Paulos CM, Carpenito C, Plesa G, Suhoski MM, Varela-Rohena A, Golovina TN, et al. The inducible costimulator (ICOS) is critical for the development of human T(H)17 cells. Sci Transl Med. 2010 Oct 27;2(55):55ra78.

79. Fos C, Salles A, Lang V, Carrette F, Audebert S, Pastor S, et al. ICOS ligation recruits the p50alpha PI3K regulatory subunit to the immunological synapse. J Immunol. 2008 Aug 1;181(3):1969–77.

80. Delgado P, Alarcón B. An orderly inactivation of intracellular retention signals controls surface expression of the T cell antigen receptor. J Exp Med. 2005 Feb 21;201(4):555–66.

81. Schutze MP, Peterson PA, Jackson MR. An N-terminal double-arginine motif maintains type II membrane proteins in the endoplasmic reticulum. The EMBO Journal. 1994 Apr 1;13(7):1696.

82. Kortleve D, Brakel M van, Wijers R, Debets R, Hammerl D. Gene Engineering T Cells with T-Cell Receptor for Adoptive Therapy. In: Immunogenetics: Methods and Protocols [Internet]. Humana; 2022.

