## Supplementary material for "Adoptive cell therapy using T cell receptors equipped with ICOS yields durable anti-tumor response": Supplenmentary information

### Supplementary information

### Supplementary Materials and Methods

*Cell culture*

Primary mouse splenocytes were isolated from spleens of Human Leukocyte Antigen (HLA)-A2.1 transgenic mice (HHD) (1,2) and were cultured at a density of $1x10^6$ cells/mL in RPMI 1640 (Gibco) with 25mM HEPES (Gibco), 10% Fetal Bovine Serum (FBS; Greiner Bio-one), Penicillin/Streptomycin (100 U/mL and 100 µg/mL, respectively, Life Technologies), 2 mM L-glutamine (Life Technologies), 1% non-essential amino acids (Gibco), 1mM sodium pyruvate (Life Technologies) and 50 µM ß-Mercaptoethanol (VWR) (Complete Mouse Medium), which was supplemented with 180 IU/mL Human Recombinant Interleukin-2 (rhIL-2) (Proleukin; Chiron). Human T cells were isolated from peripheral blood mononuclear cells (PBMCs) from healthy human donors (Sanquin) by centrifugation via Ficoll-Isopaque (density = 1.077 g/cm3; Amersham Pharmacia Biotech) and were cultured at a density of $0.5\text{-}1x10^6$ cells/mL in RPMI 1640 with 25 mM HEPES, 6% Human Serum (Sanquin), Penicillin/Streptomycin, 2 mM L-glutamine (T cell medium), which was supplemented with 360 IU/mL rhIL-2. Engineered T cells used in experiments measuring killing capacity were cultured in X-Vivo medium (Gibco), supplemented with 2% Human Serum, 110 IU/mL hIL15 (Miltenyi Biotec) and 0.1 IU/mL hIL21 (Miltenyi Biotec). The packaging cell lines Phoenix-Ampho (Ph-A, RRID: CRL-3213, kindly provided by Dr. G. Nolan, Stanford University (3)) and 293T (RRID: CRL-3216 , kindly provided by Dr. Y. Soneoka, Oxford University, Oxford, U.K. (4)) were cultured in DMEM (Gibco) with 10% FBS, 1% non-essential amino acids, 2 mM L-glutamine and Penicillin/Streptomycin (DMEM Complete). The human melanoma cell line BLM (RRID:CVCL_7035, which is HLA-A2 positive) and a BLM-gp100 clone (overexpressing the gp100 protein with or without eGFP) as well as the mouse melanoma cell lines C57BL/6 B16-F10 (ATCC CRL-6475), B16:HHD (expressing a chimeric HLA-A2/$H2K^b$ molecule (1,2)) and the B16:HHD-YLE clone (expressing a fusion protein between the human $gp100_{280\text{-}288}$ epitope YLEPGPVTA and the HHD molecule, (1)) were cultured in DMEM complete, which in case of BLM-gp100 (with or without eGFP), B16:HHD and B16:HHD-YLE was supplemented with

1mg/mL G418 (Calbiochem). The triple negative breast cancer cell line MDA-MB-231 (MM-231, RRID:HTB-26, kindly provided by Prof. J. Martens, Erasmus MC, Rotterdam, NL, which is HLA-A2 positive) and its clones overexpressing either the NY-ESO1 or the ROPN1 proteins coupled to eGFP were cultured in RPMI 1640 with 10% FBS, Penicillin/Streptomycin and 2 mM L-glutamine; in case of the MM-231-ROPN1-eGFP or MM-231-NY-ESO1-eGFP clones medium was supplemented with 2 µg/mL puromycin (Life Technologies). The TAP-deficient lymphoblastic T2 cells (RRID:CVCL_2211, obtained from ATCC: CRL1992) were cultured in RPMI 1640 with 10% FBS, 2 mM L-glutamine and Penicillin/Streptomycin. Co-culture experiments between target cells and engineered T cells were performed in RPMI 1640 with 25 mM HEPES, 10% FBS, 2 mM L-glutamine and Penicillin/Streptomycin (Cytotox medium); only in case engineered T cells were used in experiments measuring killing capacity culturing took place in X-Vivo medium (Gibco), supplemented with 2% Human Serum, 110 IU/mL hIL15 (Miltenyi Biotec) and 0.1 IU/mL hIL21 (Miltenyi Biotec). All cells were cultured at 37°C in a 5% $CO_2$ incubator and were negative for *Mycoplasma* contamination as tested on a six-weeks basis by PCR.

*Flow cytometry*

To quantify circulating TCR+ T cells in mice, blood cells were washed once with Phosphate Buffered Saline (PBS; Lonza) and incubated with 10µL of an antibody cocktail for 15 min at RT. For this, the following antibodies were used: anti-TCR-Vß14-FITC (clone CAS1.1.3, Beckman Coulter, diluted 1:2), anti-mCD3-PerCP (clone 145-2C11, BD, diluted 1:10), anti-mCD8-APC (clone 53-6.7, eBioscience, diluted 1:50) and anti-mCD4-BV650 (clone RM4-5, BD, diluted 1:2000). Following incubation, the cells were resuspended in 1% paraformaldehyde (PFA, Brunschwig) containing Flow-Count Fluorospheres (Beckman Coulter). To assess frequencies of T cell subsets, three panels of antibodies were used: (1) a T cell maturation panel consisting of: anti-mCD62L-PE (clone MEL-14, eBioscience, diluted 1:20) and anti-mCD44-APC (clone IM7, BD, diluted 1:20); (2) a T cell co-stimulation panel consisting of: anti-m4-1BB-BV421 (clone 1AH2, BD, diluted 1:20), anti-mOX40-APC (clone

OX-86, Biolegend, diluted 1:100), anti-mCD40L-PE (clone MR1, eBiosciences, diluted 1:10) and anti-mICOS-PE-Cy7 (clone 17G9, eBiosciences, diluted 1:100); and (3) a T cell co-inhibition panel consisting of: anti-mPD-1-PE-Cy7 (clone RMP1-30, Biolegend, diluted 1:100), anti-mTIM3-APC (clone B8.2C12, Biolegend, diluted 1:100), anti-mLAG3- BV421 (clone C9B7W, BD, diluted 1:20) and anti-mCTLA-4-PE (clone UC10-4B9, eBiosciences, diluted 1:20). Backbone compounds/antibodies were: viability marker 7-Amino-Actinomycin D (7-AAD; BD Biosciences, diluted 1:20), anti-mCD14-PerCP (clone rmC5-3, eBiosciences, diluted 1:20), anti-mCD3-BV510 (clone 17A2, BD Biosciences, diluted 1:100), anti-mCD4-BV650 (clone RM4-5, BD, diluted 1:2000), anti-mCD8-APC-Cy7 (clone 53-6.7, BD, diluted 1:1600) and anti-TCRVß14-FITC (clone CAS1.1.3, Beckman Coulter, diluted 1:2). To detect phosphorylated (p)PI3K, T cells were stimulated with tumor cells, fixed for 12 min at 37°C (according to Cytofix kit, BD), permeabilized for 20 min at RT, washed twice and incubated with anti-pPI3K (PE, clone PI3KY458-1A11, ThermoFisher, diluted 1:20), anti-mCD3-BV510, anti-TCRVß-14-FITC and anti-mCD8-APC-Cy7 for 30 min at RT. To detect pERK1/2, T cells were first stained with a fixable viability dye (eFluor 666, eBioscience, diluted 1:1000) for 10 min at RT, after which cells were stimulated (as described under detection of intracellular phosphorylation events), fixed for 10 min at 37°C (according to FoxP3 kit, eBioscience), washed and stained with anti-pERK1/2-FITC (clone 6B8B69, BioLegend, diluted 1:20), anti-mCD3-BV510 (clone 17A2, BD, diluted 1:100), anti-mCD8-APC-Cy7 (clone 53-6.7, BD, diluted 1:1600) and antiTCRVß14-PE (clone CAS1.1.3, Beckman Coulter, diluted 1:3) for 20 min at 4°C. To assess T cell exhaustion upon repeated stimulations with tumor cells, cells were stained with anti-mCD8, anti-mCD3, anti-TCRVß-14, anti-mPD-1 and anti-mTIM-3. To detect surface expression of hTCR:ICOS variants on human T cells, samples were stained with 7-AAD, anti-hCD3-BV421 (clone SP34-2, BD, diluted 1:200), anti-hCD8-BV650 (clone RPA-T8, BD, diluted 1:1000) and either anti-TCRVß-14-PE (gp100/A2 TCR; clone CAS1.1.3, Beckman Coulter, diluted 1:3) or anti-TCRVß-13.1-FITC (ROPN1/A2 TCR and NY-ESO1/A2 TCR; clone IMMU 222, diluted 1:10). Lastly, to assess human T cell exhaustion upon repeated stimulations, cells were stained with anti-hCD8, anti-hCD3, anti-hTCR, anti-hPD-1(APC-Cy7, clone EH12.2H7, Biolegend, diluted 1:50) and anti-hTIM-3 (APC, clone F38-2E2, Biolegend, diluted 1:50). Following above stainings, samples were washed once again, resuspended in 1% PFA, and data was acquired either with a FACS Celesta, a FACS Fortessa or a FACS Symphony A1 (BD). All flow cytometry data was analyzed using FlowJo (Treestar, version 8).

*RNA transcriptomics*

Tumor tissues were disrupted by sonification while kept on ice, and RNA was isolated (NucleoSpin, Machery Nagel). RNA concentration and quality was measured with TapeStation (Agilent), and only samples with an RNA integrity number of 6 or higher and a quantity of more than 250 ng were used for 3' RNA sequencing (QuantSeq, Illumina, Erasmus MC). Sequences were mapped to mouse reference genome mm10. Differential gene expression analysis was performed using the DESeq2 package from R, and gene set enrichment was performed with the GSA and fgsea packages (5–7). Data was filtered to keep only genes that have a higher overall count of >10 across all samples. References used in gene set enrichment analyses were a laboratory list of immune-related genes that are associated to immune escape (8), and the mouse Hallmark database (available at "https://data.broadinstitute.org/gsea-msigdb/msigdb/release/2022.1.Mm/ ", mh.all.v2022.1.Mm.symbols.gmt).

### References


1. Straetemans T, Berrevoets C, Coccoris M, Treffers-Westerlaken E, Wijers R, Cole DK, et al. Recurrence of Melanoma Following T Cell Treatment: Continued Antigen Expression in a Tumor That Evades T Cell Recruitment. Mol Ther. 2015 Feb;23(2):396–406.

2. Schaft N, Willemsen RA, de Vries J, Lankiewicz B, Essers BWL, Gratama JW, et al. Peptide fine specificity of anti-glycoprotein 100 CTL is preserved following transfer of engineered TCR alpha beta genes into primary human T lymphocytes. J Immunol. 2003 Feb 15;170(4):2186–94.

3. Weijtens ME, Willemsen RA, Hart EH, Bolhuis RL. A retroviral vector system 'STITCH' in combination with an optimized single chain antibody chimeric receptor gene structure allows efficient gene transduction and expression in human T lymphocytes. Gene Ther. 1998 Sep;5(9):1195–203.

4. Grignani F, Kinsella T, Mencarelli A, Valtieri M, Riganelli D, Grignani F, et al. High-efficiency gene transfer and selection of human hematopoietic progenitor cells with a hybrid EBV/retroviral vector expressing the green fluorescence protein. Cancer Res. 1998 Jan 1;58(1):14–9.

5. Love MI, Huber W, Anders S. Moderated estimation of fold change and dispersion for RNA-seq data with DESeq2. Genome Biology. 2014 Dec 5;15(12):550.

6. Subramanian A, Tamayo P, Mootha VK, Mukherjee S, Ebert BL, Gillette MA, et al. Gene set enrichment analysis: a knowledge-based approach for interpreting genome-wide expression profiles. Proc Natl Acad Sci U S A. 2005 Oct 25;102(43):15545–50.

7. Korotkevich G, Sukhov V, Budin N, Shpak B, Artyomov MN, Sergushichev A. Fast gene set enrichment analysis [Internet]. bioRxiv; 2021 [cited 2024 Dec 8]. p. 060012. Available from: https://www.biorxiv.org/content/10.1101/060012v3

8. Hammerl D, Massink MPG, Smid M, van Deurzen CHM, Meijers-Heijboer HEJ, Waisfisz Q, et al. Clonality, Antigen Recognition, and Suppression of CD8+ T Cells Differentially Affect Prognosis of Breast Cancer Subtypes. Clin Cancer Res. 2020 Jan 15;26(2):505–17.

### Supplementary Figure 1

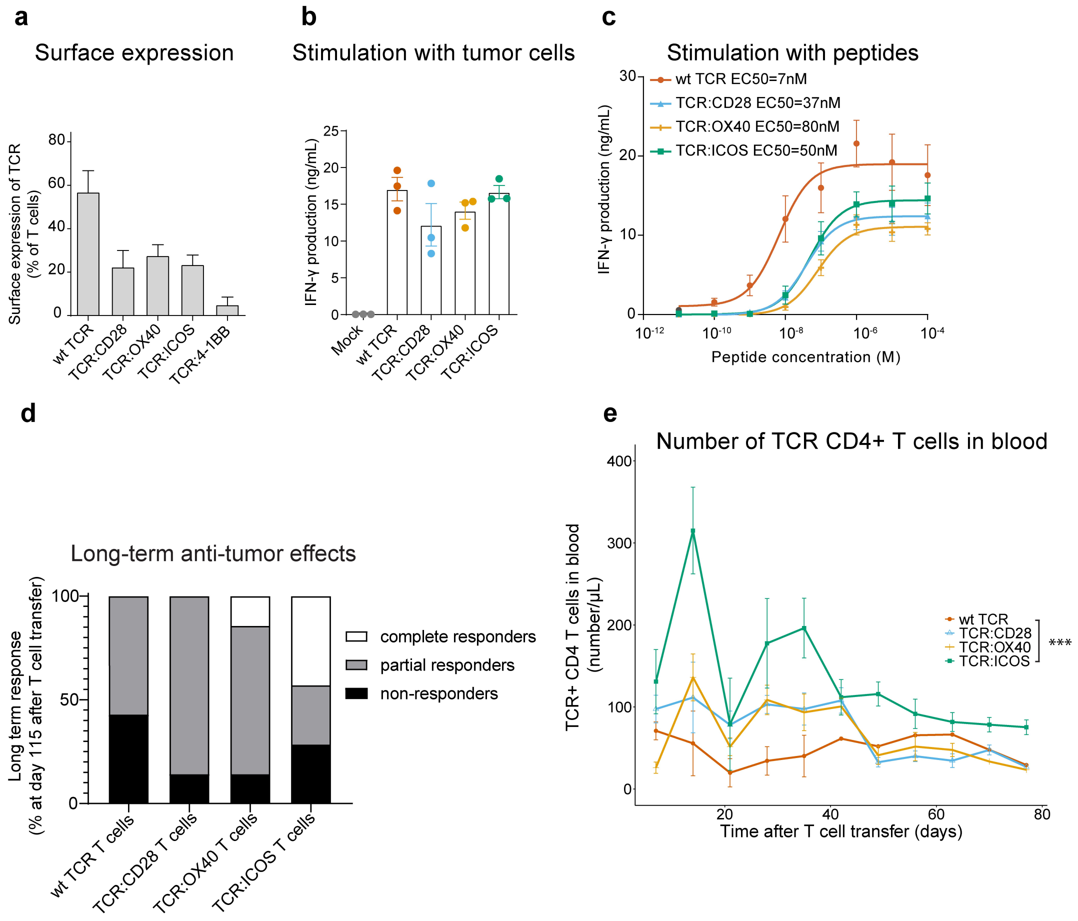


***Supplementary Figure 1.*** *In vitro and in vivo performance of mouse T cells expressing co-stimulatory TCRs.* **(a)** Bar graph showing surface expression of co-stimulatory TCRs after transduction of mouse splenocytes (mean±SEM, n=2-4). **(b)** Bar graph showing IFN-γ production of TCR T cells following overnight stimulation with B16-HHD-YLE tumor cells (mean±SEM, n=3). **(c)** IFN-γ production is plotted following stimulation with B16-HHD cells that were pre-loaded with titrated amounts of human gp100 peptide. TCR T cells were co-cultured with peptide-loaded T2 cells for 16h at 37°C and supernatants were assayed using an IFN-γ ELISA (mean±SEM, n=6). EC50 values were calculated using graphpad prism. **(d)** Stacked bars displaying overall anti-tumor response rate of mice treated with different TCR T cells (fraction of total, n=7). Definitions of complete, partial and non-response are given in Materials and Methods section. **(e)** Line graph showing number of TCR+ CD4 T cells per µL blood following adoptive T cell transfer in melanoma mouse model as determined by flow cytometry (median±SEM, n=4-6). Inter-group comparison was done using a linear mixed model with elapsed time as a separate variable and correction for random effect. With *** = p<0.001.

### Supplementary Figure 2

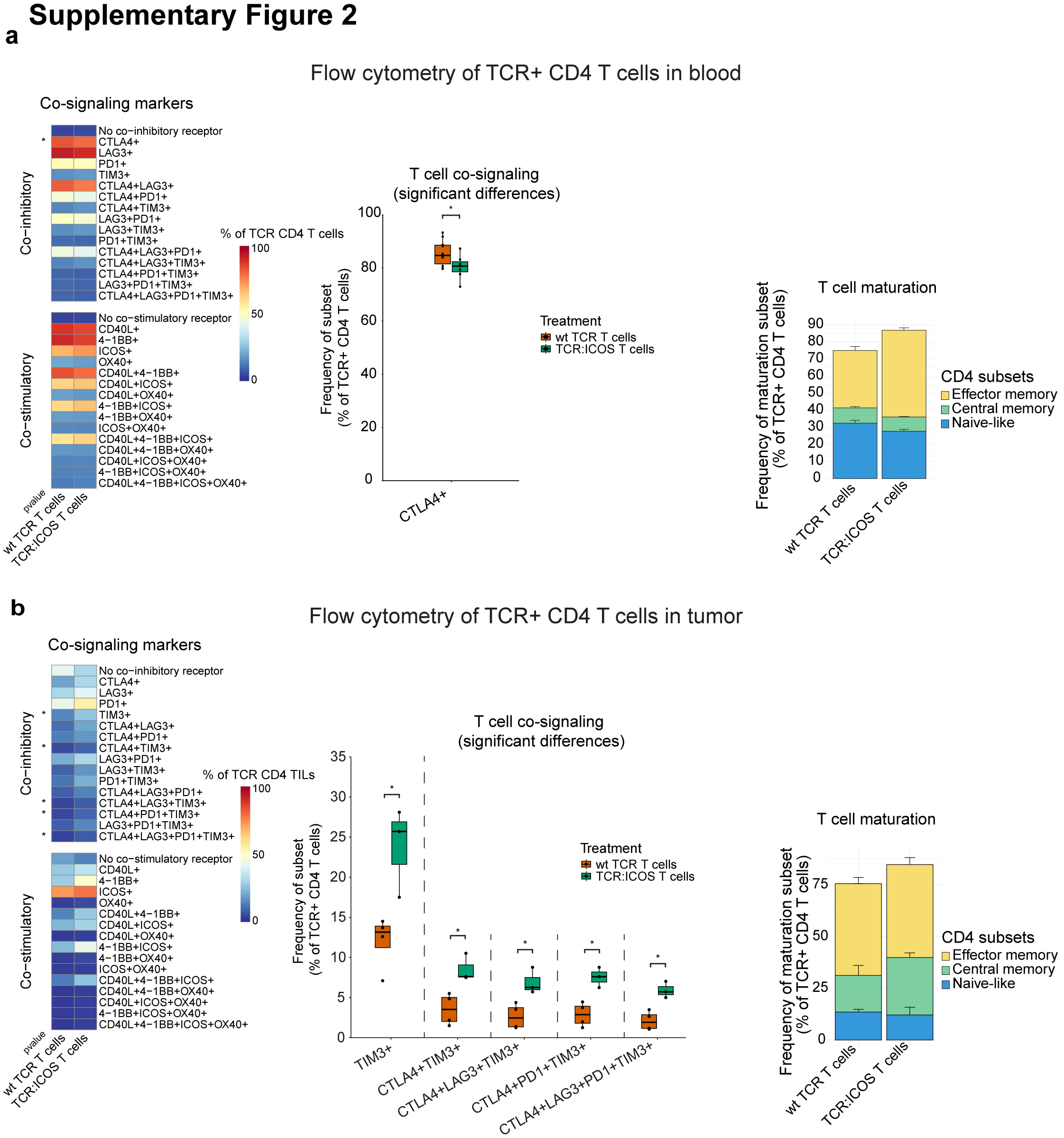


***Supplementary Figure 2.*** *Phenotypic analysis of TCR+ CD4 T cells upon adoptive transfer of TCR:ICOS T cells in melanoma mouse model.* ***(a)*** Left panel: heatmap corresponding to phenotype of TCR+ CD4 T cells in blood three weeks after T cell transfer in melanoma mouse model as determined by flow cytometry. Co-signaling phenotype of T cells was determined according to the expression of (combinations of) co-inhibitory (CTLA4, LAG3, PD1, TIM3) or co-stimulatory receptors (CD40L, CD137, ICOS, OX40). The heatmap displays median expressions of single or multiple markers, n=3-8 per group. Statistical significance between TCR:ICOS and wt TCR T cells was calculated using a t-test test and is highlighted on the left side of the heatmap. Middle panel: TCR:ICOS+ CD4 T cell subsets that show differential frequencies when compared to wt TCR+ CD4 T cells. Right panel: stacked bars corresponding to maturation phenotype of TCR+ CD4 T cells in blood as determined by flow cytometry (n=3-8 per group). Maturation status of T cells was defined as follows: naïve T cells, CD62L+CD44; central memory T cells, CD62L+CD44+; and effector memory T cells, CD62L-CD44+. **(b)** Left and right panels: heatmap and stacked bars corresponding to phenotype of TCR+ CD4 TILs in regressing tumors at five days after T cell transfer as determined by flow cytometry (n=2-4 per group). Statistical testing of TCR:ICOS CD4 TIL subsets was not always possible due to too low number of cells detected. Middle panel: TCR:ICOS+ CD4 TIL subsets that show differential frequencies when compared to wt TCR+ CD4 TILs. With * = $p<0.05$.

### Supplementary Figure 3

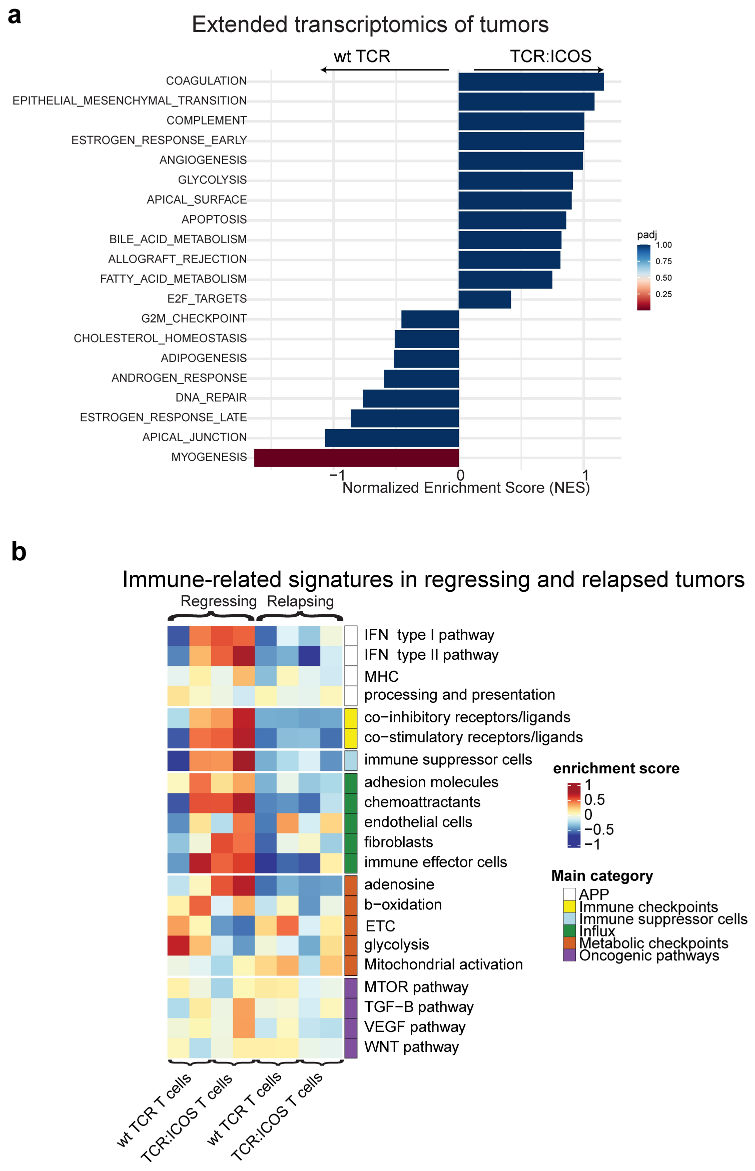


***Supplementary Figure 3.*** *Transcriptomic analysis of relapsing tumors upon adoptive transfer of TCR:ICOS T cells in melanoma mouse model.* Relapsing B16-HHD-YLE tumors of mice treated with adoptive cell treatment using TCR:ICOS or wt TCR T cells were isolated, and their transcriptome was analyzed. **(a)** Barplot showing top 20 gene sets that are enriched in relapsing tumors following transfer of TCR:ICOS T cells versus wt TCR T cells. **(b)** Heatmap displaying enrichment of immune-related signatures that are associated with immune-escape in both regressed and relapsed tumors from wt TCR and TCR:ICOS T cell-treated mice.

### Supplementary Figure 4

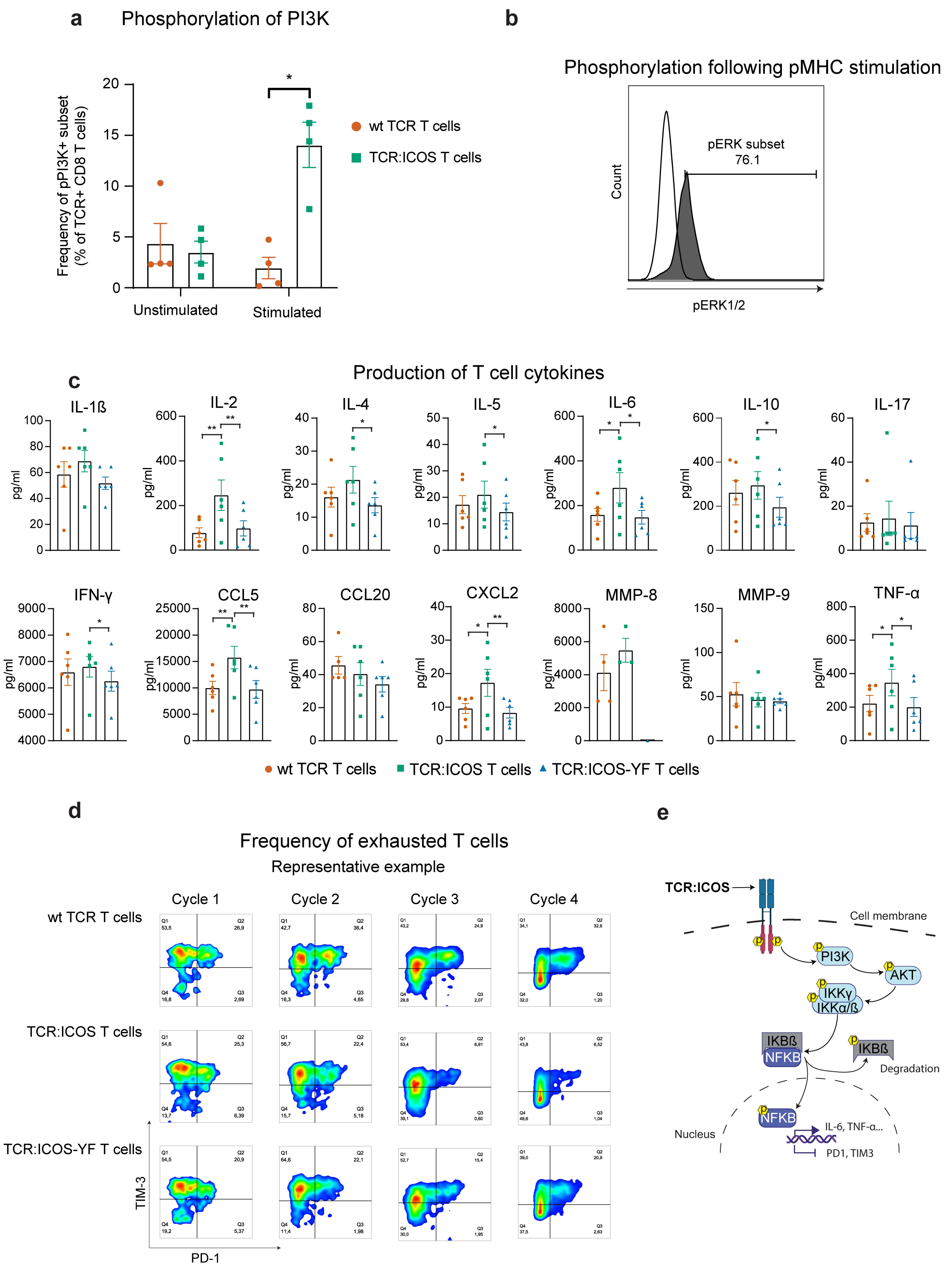

***Supplementary Figure 4.*** *In vitro characterization of TCR:ICOS T cells with example data and proposed mechanism of action.* **(a)** Bar graph displaying phosphorylation of PI3K following stimulation of mouse T cells with tumor cells. T cells were incubated for 2h at 37°C with B16F10 cells that either expressed the cognate epitope or not, after which T cells were stained and analyzed for pPI3K using flow cytometry (n=4). Statistical significance between TCR:ICOS and wt TCR T cells was tested using a two-way ANOVA with Sidack's multiple comparison test. **(b)** Validation of the stimulation conditions used for the phosphorylation array in Figure 4B, tested by flow cytometric detection of the phosphorylation of ERK1/2. Wildtype TCR T cells were stimulated with either cognate peptide (grey filled curve) or irrelevant peptide (white filled curve) for 30 min at 37°C, before cells were stained for pERK1/2 and measured using flow cytometry. **(c)** Bar graphs displaying cytokine production following stimulation with tumor cells. T cells were incubated for 16h at 37°C with B16 cells that either expressed the cognate epitope or not, after which supernatants were assayed for the presence of 14 cytokines according to a cytokine bead array (mean±SEM, n=1-6). Statistical significance between TCR:ICOS and wt TCR T cells was tested with a two-way ANOVA with Dunnet's multiple comparison test. **(d)** Representative example of flow cytometric changes in % of $PD1^{+}TIM3^{+}$ TCR+ CD8 T cells following repeated antigen stimulations. **(e)** Schematic illustration of proposed mechanism of action of TCR:ICOS. With * = $p<0.05$, **=$p<0.01$, ***=$p<0.005$

### Supplementary Figure 5

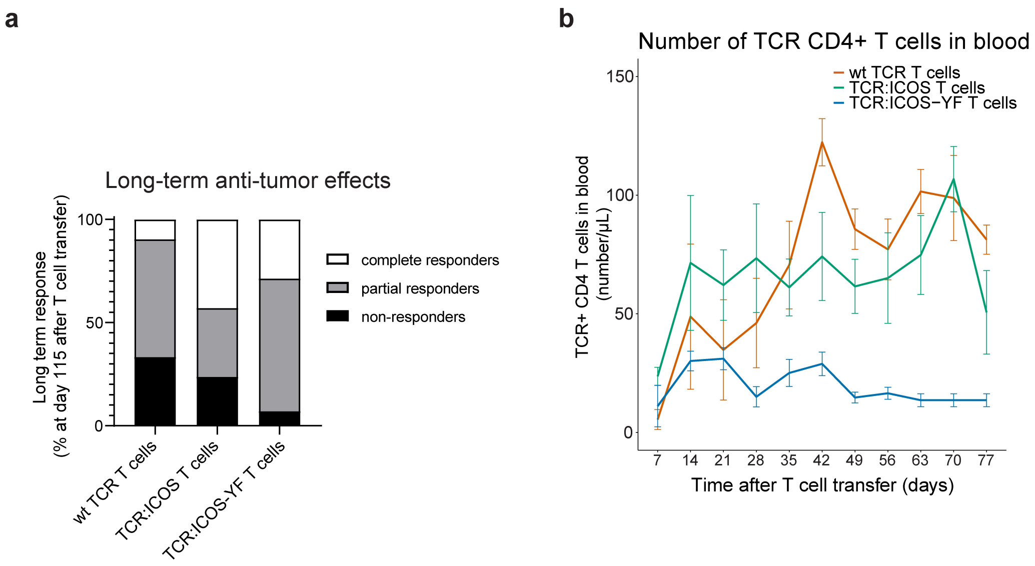


***Supplementary Figure 5.*** *In vivo anti-tumor response of TCR:ICOS-YF T cells upon adoptive transfer in melanoma mouse model.* **(a)** Stacked bars displaying overall anti-tumor response rate of mice treated with different TCR T cells (fraction of total, n=6). Definitions of complete, partial and non-response are given in Materials and Methods section. **(b)** Numbers of TCR+ CD4 T cells per µL blood following T cell transfer in melanoma mouse model is plotted as determined by flow cytometry (median±SEM, n=4-6). Inter-group comparison was done using a linear mixed model test with elapsed time as a separate variable and correction for random effect.

### Supplementary Figure 6

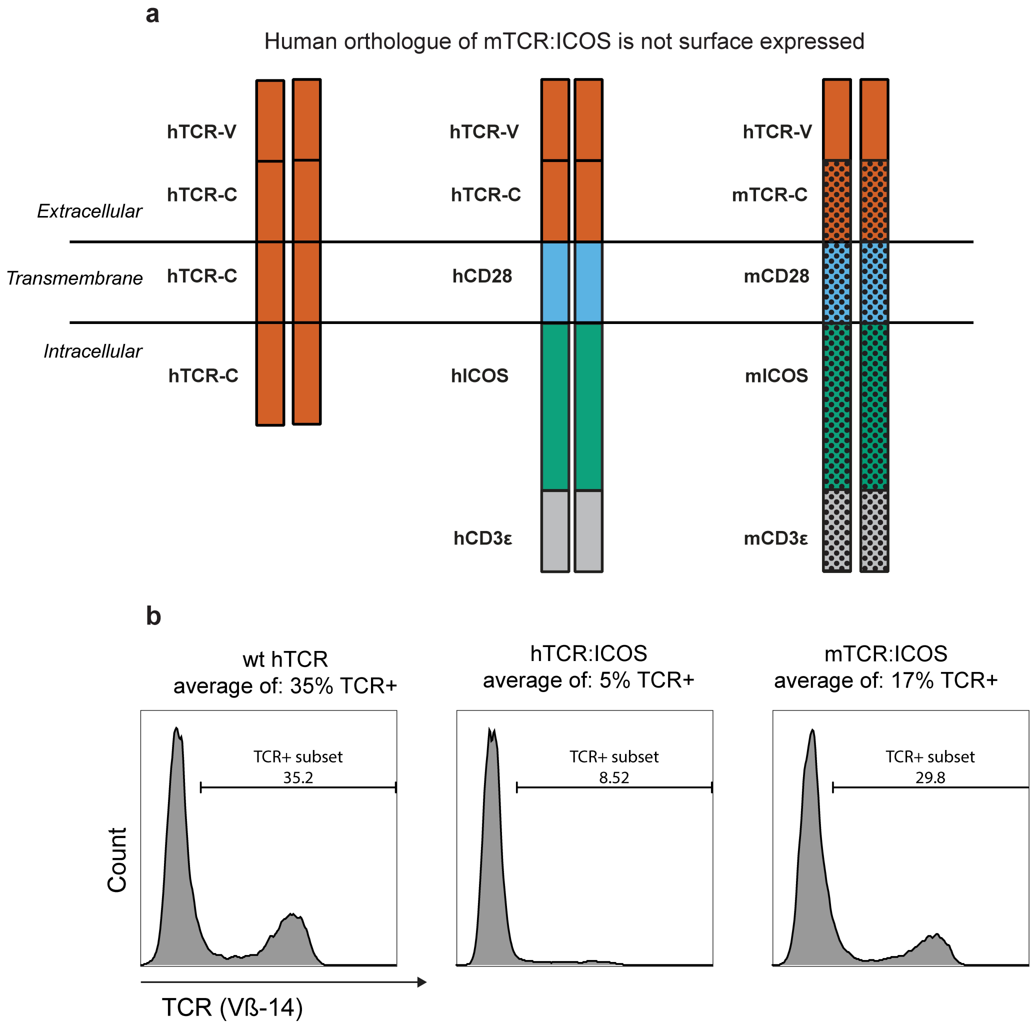


***Supplementary Figure 6.*** *Surface expression of human orthologue of mTCR:ICOS on human T cells.* **(a)** Top panel: graphical representation of wt TCR, mouse (m) TCR:ICOS and human (h) TCR:ICOS transgenes, in which mouse domains are depicted with dots. **(b)** Surface expression in human T cells according to % TCR-Vß14+ cells within all CD3+ cells (mean, n=32, 18, 24 for wt TCR, hTCR:ICOS and mTCR:ICOS respectively) and example of flow cytometric histograms displaying expression of the different TCRs on human primary T cells.
